# Inoculation of *Malus baccata* ‘Jackii’-derived offspring and QTL analysis reveal a polygenic inheritance pattern of apple blotch resistance

**DOI:** 10.64898/2026.04.09.717374

**Authors:** Matthias Pfeifer, Andreas Peil, Henryk Flachowsky, Ofere Francis Emeriewen, Thomas Wolfgang Wöhner

## Abstract

Apple blotch, caused by *Diplocarpon coronariae*, is an increasingly important fungal disease that leads to premature leaf fall and significant yield losses in apple orchards. Breeding resistant cultivars offers a sustainable strategy to reduce disease impact, as all commercial apple cultivars are susceptible to this pathogen. This study aimed to investigate the disease resistance of *Malus baccata* ‘Jackii’-derived offspring to *D. coronariae* through artificial inoculation and to identify loci associated with resistance. Simple interval mapping was performed using phenotypic and genotypic data from 122 individuals of an F_1_ population (’Idared’ × *M. baccata* ‘Jackii’), together with analyses of *M. baccata* ‘Jackii’-derived open-pollinated populations. Our results indicate that resistance to apple blotch is a complex, polygenic trait, with four important QTLs identified on linkage groups 1, 2, 12 and 13. Disease severity was strongly affected by inoculum, phenotyping method and environmental factors. These findings have direct implications for apple breeding programmes aimed at developing apple blotch-resistant cultivars.

## Introduction

*Diplocarpon coronariae* (Ellis & Davis) Wöhner & Rossman is the causal agent of apple blotch, also known as premature leaf fall (Wöhner and Emeriewen, 2019; Crous et al., 2020). The pathogen had already been described in North America (Davis, 1903) and Japan (Engler, 1906) at the beginning of the 20th century. Since the late 20^th^ century, apple blotch has become increasingly problematic in Asian countries such as India (Sharma et al., 2004) and Korea (Kim et al., 1998), where it is now considered one of the most important apple diseases (Lee et al., 2006; Kwon et al., 2015; Watpade et al., 2021). More recently, the disease has also gained importance in the USA (Khodadadi et al., 2022) and Europe (Wöhner and Emeriewen, 2019).

*D. coronariae* is a heterothallic ascomycete (Cheng et al., 2021) capable of producing both conidia and ascospores (Wöhner and Emeriewen, 2019). While the role of ascospores in the disease cycle is not yet fully understood, conidia can multiply rapidly, leading to multiple infection cycles within a single growing season (Zhao et al., 2013; Wöhner and Emeriewen, 2019). Conidial germination typically occurs within a few hours of leaf wetness, depending on temperature (Lian et al., 2021). After penetration of the cuticle, haustoria and hyphae are formed, and at approximately seven days post-inoculation (dpi), acervuli develop (Zhao et al., 2013). The fungus forms haustoria as well as subcuticular and intercellular hyphae, followed later by intracellular hyphae and subcuticular hyphal strands, and it has been suggested that it may exhibit a hemibiotrophic infection strategy (Zhao et al., 2013). The disease is dispersed by wind and splash mechanisms and its spread has been associated with rainfall events (Dong et al., 2015; Kim et al., 2019; Boutry et al., 2023), consistent with the requirement of leaf wetness for extensive infection (Lian et al., 2021). Typical symptoms include dark spots on the adaxial leaf surface, which develop into thread-like structures, causing chlorosis and ultimately premature leaf fall, weakening the tree and leading to significant yield loss (Wöhner and Emeriewen, 2019).

Although effective disease management is possible through plant protection measures (Sharma et al., 2004; Dang et al., 2017; Watpade et al., 2021; Khodadadi et al., 2022), cultivation of resistant or tolerant cultivars represents a sustainable strategy to reduce yield losses. However, no immune apple cultivars are currently known (Richter et al., 2025a; Wöhner et al., 2021). Resistance in commercial apple cultivars is often limited, with differences in leaf fall spanning only a few days (Khodadadi et al., 2022). Therefore, the identification of resistance or tolerance in wild *Malus* species is of great interest. *Malus baccata* ‘Jackii’ (accession number: MAL0419; hereafter referred to as *Mb*j) has been reported to exhibit resistance to *D. coronariae* (Wöhner et al., 2021; Pfeifer et al., 2024). This genotype is also resistant to powdery mildew (Dunemann and Schuster, 2009; Pfeifer et al., 2026a), apple scab (Gygax et al., 2004), fire blight (Vogt et al., 2013; Wöhner et al., 2016; Wöhner et al., 2018) and potentially to some fruit rot pathogens (Pfeifer et al., 2025), making it a particularly valuable resource for resistance breeding. However, the genetic basis of *Mb*j-derived resistance to *D. coronariae* has so far remained unclear and therefore cannot be efficiently exploited in breeding programmes. In this study, we aimed to dissect the genetic architecture of *Mb*j-derived resistance to *D. coronariae*. To this end, three main objectives were pursued. First, we tested whether the trait is controlled by a major monogenic dominant resistance factor through artificial inoculation of F_1_ progeny (’Idared’ × *Mb*j) combined with quantitative trait locus (QTL) analysis. Subsequently, the temporal dynamics of symptom development were characterised and integrated with genotypic data to identify QTLs associated with disease progression. Finally, additional open-pollinated populations derived from *Mb*j and two F_1_ individuals were evaluated to assess the potential contribution of a major recessive locus.

## Materials and methods

### Plant material

To analyse the segregation of resistance and to assess its mode of inheritance, a biparental F_1_ population (’Idared’ × *Mb*j) was generated. A total of 122 F_1_ individuals obtained from the cross ’Idared’ × *Mb*j (populations 05225 and 06228), including both parents, were cultivated in the experimental orchard of the Julius Kühn-Institut (JKI) in Dresden-Pillnitz, Germany, without fungicide protection. The F_1_ population and the parental genotypes were additionally maintained under greenhouse conditions and at a second outdoor site as potted trees. To complement the mapping F_1_ population, three additional populations derived from open pollination were generated at the second outdoor site. These populations were established to further examine the inheritance pattern of resistance, particularly to assess the possibility of a recessive genetic component. Situated approximately 220 m from the experimental orchard, the second outdoor site was considered to favour predominantly within-population pollination. Therefore, seeds were collected in October 2023 from *Mb*j and the two F_1_ individuals 05225-31 and 06228-66. In total, 232 *Mb*j-derived seedlings (‘Jackii’-OA), 40 seedlings from 05225-31 (05225-31-OA), and 30 seedlings from 06228-66 (06228-66-OA) were grown under greenhouse conditions and analysed.

### Genotyping and genetic linkage map construction of *Mb*j

Genotyping of the F_1_ population and construction of the genetic linkage map using JoinMap 5 (van Ooijen, 2018) are described in detail in previous studies (Pfeifer et al., 2026a; Pfeifer et al., 2026b). In short, all F_1_ individuals and both parental genotypes were genotyped using tunable genotyping-by-sequencing (Ott et al., 2017) with the restriction enzyme *Bsp*1286I. Only quality-filtered single nucleotide polymorphism (SNP) sequences that uniquely aligned to *Mb*j haplotype 1 were retained for linkage map construction. In addition, 70 simple sequence repeat (SSR) markers were used for genotyping. These markers were amplified by multiplex PCR using the Type-it Microsatellite PCR Kit (Qiagen, Hilden, Germany), analysed on an Applied Biosystems 3500xL Genetic Analyzer and processed using GeneMapper v6 (Applied Biosystems, Waltham, MA, USA). SSR markers were subsequently integrated into the genetic linkage map, which was constructed using a regression mapping algorithm and Kosambi’s mapping function.

### SSR genotyping of open-pollinated progeny and removal of undesired offspring

Open-pollinated genotypes were screened using established SSR markers to exclude undesired crosses with unrelated genotypes, the parental genotypes ’Idared’ and *Mb*j, and self-fertilisation, thereby retaining only progeny derived from pollen donors originating from the F_1_ population (’Idared’ × *Mb*j). DNA was extracted from young leaves using the REDExtract-N-Amp Plant PCR-Kit (Sigma-Aldrich, St. Louis, MO, USA), and PCR was performed with the Type-it Microsatellite PCR Kit (Qiagen, Hilden, Germany) employing 11 established SSR markers (Table S1). The primers were divided into two multiplexes, with forward primers labelled with different fluorescent dyes, and fragments were assigned to the primers based on expected fragment sizes reported in the literature (Hokanson et al., 1998; Liebhard et al., 2002). The PCR mixture contained 1 μl of primer mix (1 pmol/µl of each primer), 2 μl of ddH_2_O, 5 μl of Type-it Multiplex PCR Master Mix, 1 μl of Q-solution, and 1 μl of DNA. PCR was performed at 95 °C for 5 min, followed by 40 cycles of 95 °C for 30 s, 55 °C for 1 min 30 s, and 72 °C for 1 min, and concluded with 60 °C for 30 min. 1 μl of the 1:100 diluted PCR product was mixed with 9 μl ABI solution (1 ml Hi-Di formamide + 6 μl GeneScan-600 LIZ size standard; Applied Biosystems, Waltham, MA, USA). After denaturation at 95 °C for 5 min, the samples were analysed on an Applied Biosystems 3500xL Genetic Analyzer, and SSR alleles were scored using GeneMapper v6 (Applied Biosystems, Waltham, MA, USA). Genotypes were classified as outcrossers and excluded from the analysis if they carried at least one allele absent in either ’Idared’ or *Mb*j and differing by ≥ 2 bp from the parental alleles. To exclude self-fertilisation, offspring were required to possess at least one allele not present in the mother. In the *Mb*j open-pollinated population, genotypes were further excluded if they consistently carried one allele from ’Idared’ across all 11 markers, as these likely represented *Mb*j × ’Idared’ offspring rather than the desired *Mb*j × F_1_ cross. Open-pollinated 05225-31- and 06228-66-derived genotypes were removed if a backcross to ’Idared’ or *Mb*j could not be ruled out. This applied to genotypes in which all alleles not present in the mother could be exclusively assigned to ’Idared’ or *Mb*j. Following this selection process, only genotypes resulting from crosses with F_1_ descendants of ’Idared’ × *Mb*j were retained.

### SSR genotyping of *D. coronariae* used in inoculation experiments

Leaves infected with *D. coronariae,* collected in 2022 and 2023 and used for inoculum preparation in this study, were genotyped with 12 SSR markers to determine multilocus genotypes (MLGs) based on Oberhänsli et al. (2021) with minor modifications. Several leaves collected in each year were frozen in liquid nitrogen, homogenised, and DNA was extracted using the DNeasy Plant Mini Kit (Qiagen, Hilden, Germany) according to the manufacturer’s protocol. DNA concentrations were adjusted to 10 ng/μl using a NanoDrop One^C^ spectrophotometer (Thermo Fisher Scientific Inc., Waltham, MA, USA). Primer pairs were multiplexed as described (Oberhänsli et al., 2021), and PCR was performed using the Type-it Microsatellite PCR Kit (Qiagen, Hilden, Germany) with 1 μl of primer mix (1 pmol/µl of each primer), 1 μl of ddH_2_O, 5 μl of Type-it Multiplex PCR Master Mix, 1 μl of Q-solution, and 2 μl of DNA. PCR was performed with 40 cycles instead of 35 (Oberhänsli et al., 2021) . Subsequently, PCR products were diluted 1:100 in ddH₂O and analysed following the same procedure as described above for the other SSR markers.

### Inoculum preparation

Infected leaves of the apple cultivar ’Topaz’ were collected in autumn 2022 and 2023 from an orchard heavily infected with apple blotch at the Competence Centre for Fruit Growing, Bavendorf, Lake Constance, Germany. The leaves were stored at -20 °C until use. A conidial suspension of *D. coronariae* was prepared by mixing frozen, infected apple leaves with water supplemented with 0.05% Tween 20 (Sigma-Aldrich, St. Louis, MO, USA). Concentration of conidia was determined using a haemocytometer (Fein-Optik, Bad Blankenburg, Germany) and a Jenaval microscope (Carl Zeiss AG, Oberkochen, Germany) and adjusted to 10^5^ conidia per ml. Conidial germination was verified with several 10 μl droplets of the suspension placed onto 1.5% water agar and incubated at room temperature for 48 h with subsequent evaluation using an Axiovert 25 microscope (Carl Zeiss AG, Oberkochen, Germany).

### Detached leaf assay and image analysis

Detached leaf assays were conducted to evaluate the inheritance of resistance to apple blotch. The F_1_ population was inoculated three times in 2023 using inoculum collected in 2022 and once in 2025 using inoculum collected in 2023. For populations derived from open pollination, the detached leaf assay was performed once in 2024 using inoculum from 2022 and twice using inoculum from 2023. For this, leaves from greenhouse- and field-grown plants were inoculated by spraying a conidial suspension with a spray bottle on the adaxial and abaxial leaf surfaces. Two inoculated leaves were placed per Petri dish (145 x 20 mm; Greiner Bio-One, Kremsmünster, Austria) on a circular filter paper (125 mm diameter) and a curved grid to prevent direct contact between the leaves and the water at the bottom of the Petri dish. A total of 7.5 ml of water was added to each Petri dish. The Petri dishes were stacked, sealed in plastic bags, and incubated at 24 °C under a 12 h light/dark cycle. Four leaves per genotype and repetition were inoculated.

Leaf images were captured at 7, 10 and 14 dpi on a Prolite scan SC light box (Kaiser Fototechnik, Buchen, Germany) using an EOS 90D camera (Canon, Tokyo, Japan). Image analysis comprised leaf area determination, background removal, calculation of the percentage of necrotic leaf area, and counting of necrotic spots. Leaf area was quantified using the Python script Leaf_area_calculator.py (Supplementary script S1), which relies on a reference square included in each image. Background removal and image resizing were performed with Leaf_extractor_resizer.py (Supplementary script S2), and the percentage of necrotic leaf area was quantified using Leaf_color_detector.py (Supplementary script S3). Apple blotch-specific necrotic spots were counted using a custom-developed YOLOv5s model (Reim et al., 2024) with a confidence threshold of 0.45 and an intersection-over-union threshold of 0.7. The model was trained to detect individual apple blotch-specific necrotic spots on healthy green leaf tissue and does not identify necrotic spots within large, contiguous brown areas. An example of a leaf before and after processing with the Python scripts and the YOLOv5s model is shown in Figure 1.

**Figure 1.**
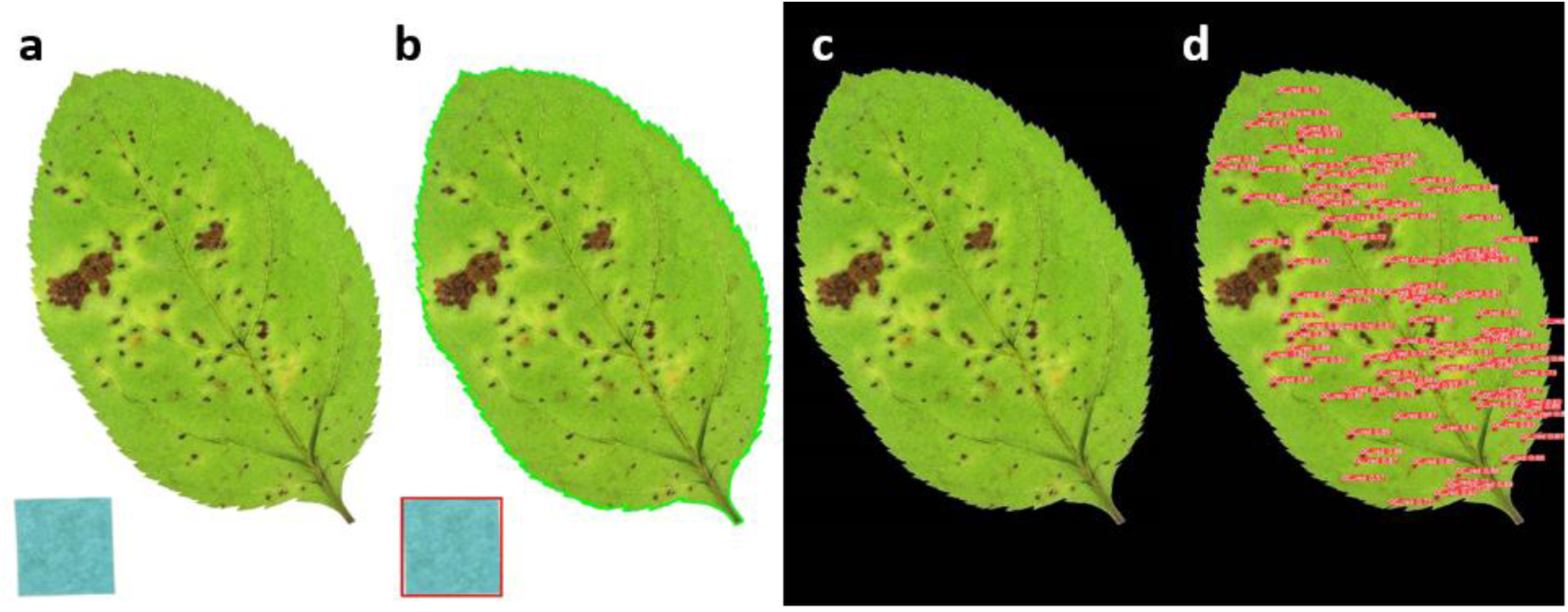
A leaf from the detached leaf assay before and after processing using the Python scripts and the YOLOv5s model: (a) original image; (b) leaf area quantification output; (c) extracted and resized leaf image; (d) leaf showing marked necrotic spots counted.

### Greenhouse experiment

In addition, artificial inoculation experiments were conducted on living plants under greenhouse conditions. Leaves from the sixth node downward were sprayed with inoculum on the adaxial and abaxial leaf surfaces. In 2023, two treatments were applied to the F_1_ population of ’Idared’ × *Mb*j: treatment with or without a zip bag. For the former, inoculated leaves were enclosed in a zip bag immediately after inoculation and kept enclosed for three days to maintain high humidity and leaf wetness (Figure 2). After three days, the bags were removed. Both treatments were applied to the same plant at the same time using the same inoculum to allow direct comparison of treatment effects. For the populations derived from open pollination and for the greenhouse experiments conducted in 2025, only the zip bag treatment was applied. Greenhouse plants were maintained at approximately 25 °C from 6 am to 8 pm and at 20 °C during the night. Automatic mist irrigation (20 s every 10 min) was used to ensure continuous leaf wetness and high humidity, thereby promoting apple blotch infection. Disease symptoms were evaluated at 7-day intervals, starting at 14 dpi and continuing until 91 or 98 dpi, using the phenotyping scale shown in Figure 3. Greenhouse experiments of the F_1_ population were conducted three times in 2023 using the inoculum from 2022 and once each in 2025 using inoculum from 2022 and 2023, respectively. In contrast, the populations derived from open pollination were inoculated once in 2024 using inoculum from 2023. For each repetition, three leaves per treatment and genotype were inoculated.

**Figure 2.**
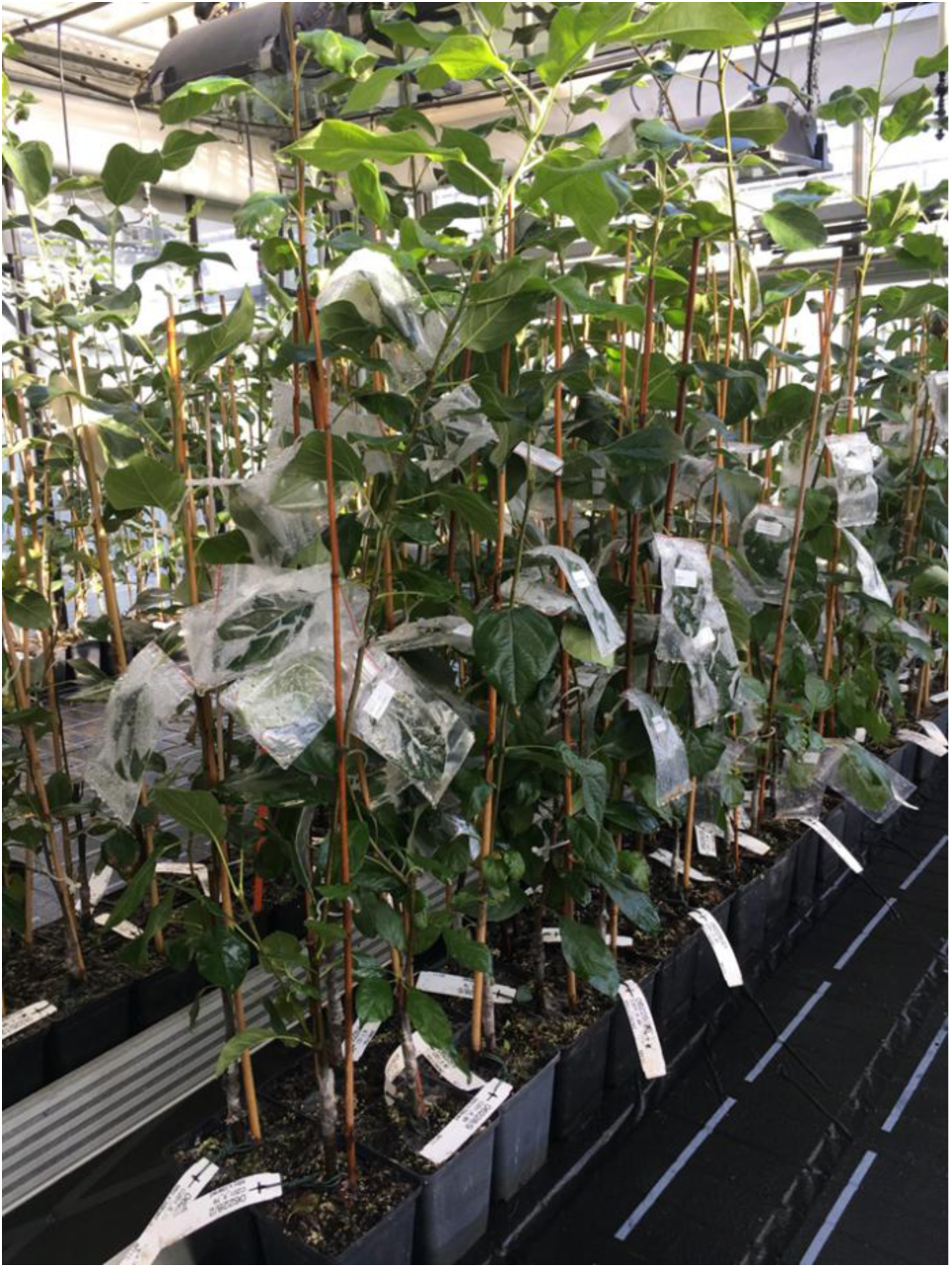
Inoculated greenhouse plants showingleaves with and without zip bag treatment on the same plant.

**Figure 3.**
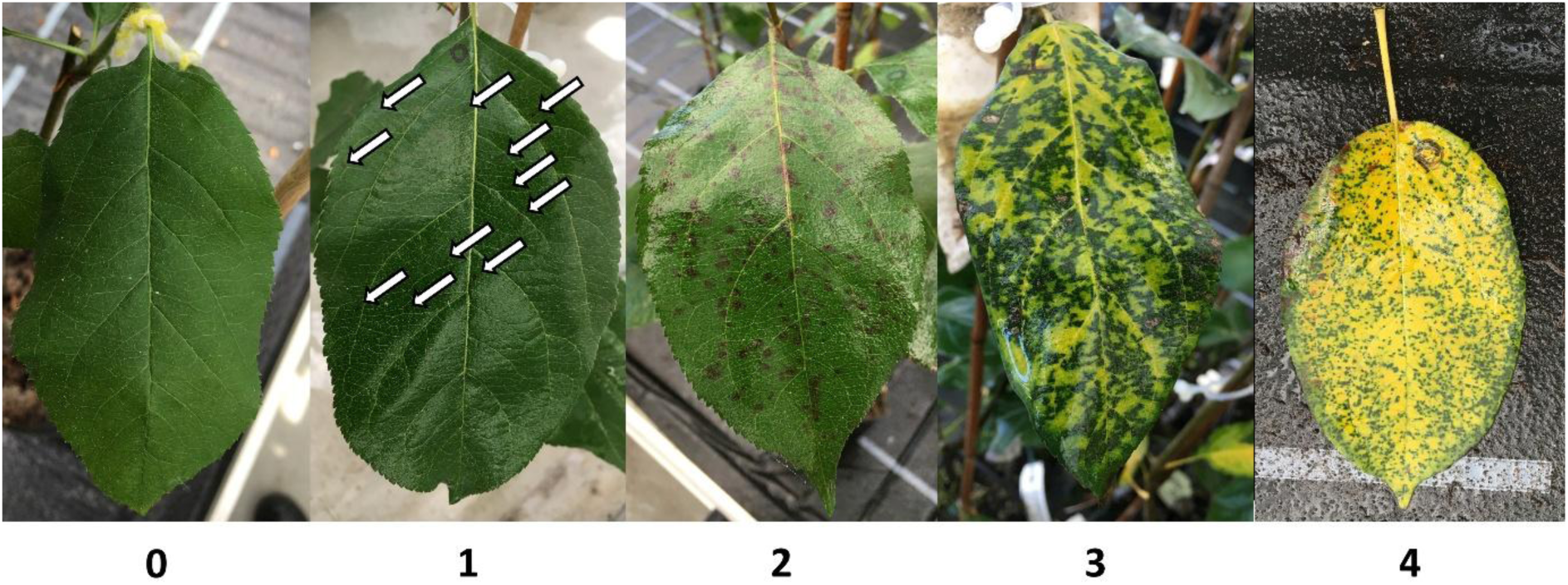
Phenotyping scale used in the greenhouse experiment: 0 = no necrotic spots; 1 = few necrotic spots; 2 = numerous necrotic spots or spreading infection; 3 = extensive necroses or chloroses; 4 = leaf drop.

### Microscopic analyses

At 21 dpi, leaves from the third repetition of the detached leaf assay (F_1_ population, 2023) were examined with a Stemi 1000 stereomicroscope (Carl Zeiss AG, Oberkochen, Germany). *D. coronariae*-specific symptoms were classified into four categories: (4a) point necrosis with an acervulus; (4b) point necrosis with an acervulus, surrounded by circular desiccated light-brown necrotic tissue; (4c) acervuli connected by visible mycelial threads; and (4d) necrosis with acervuli surrounded by dark-brown tissue (Figure 4). The total number of acervuli was counted on five necrotic lesions per genotype.

**Figure 4.**
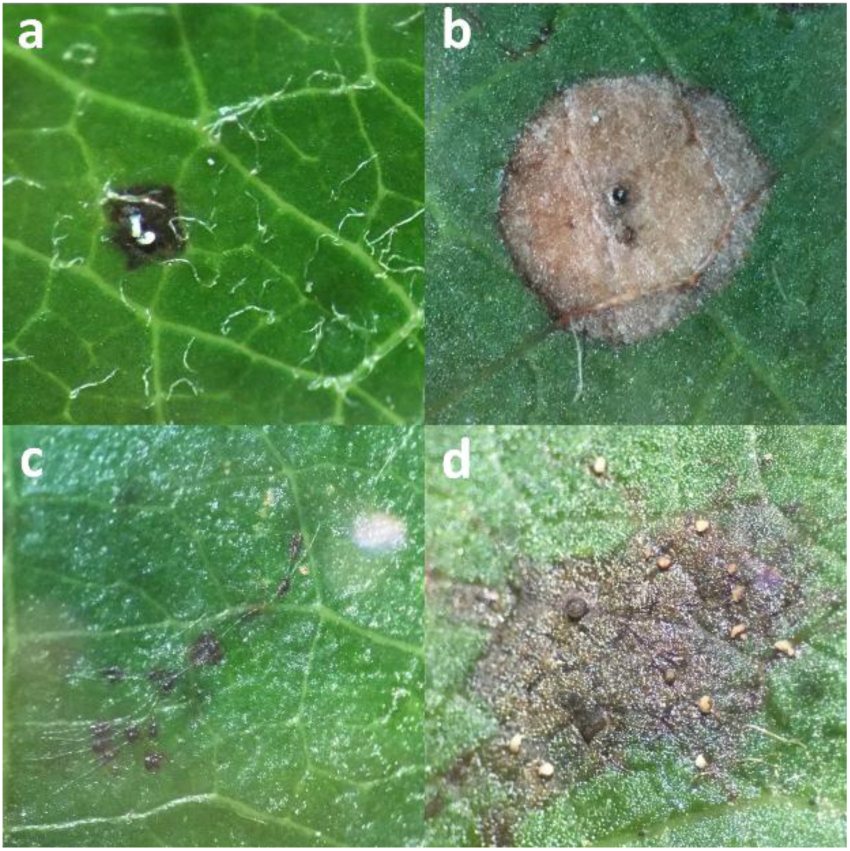
Different symptoms of *D. coronariae* observed in the detached leaf assay on apple leaves: (a) point necrosis with an acervulus; (b) point necrosis with an acervulus, surrounded by circular desiccated light-brown necrotic tissue; (c) acervuli connected by visible mycelial threads; (d) necrosis with acervuli surrounded by dark-brown tissue.

For detailed analysis, leaves from *Mb*j were examined further. Leaf discs originating from both the detached leaf assay and the greenhouse experiment were destained in ethanol-acetic acid (3:1) for ≥ 24 h. Afterremoval of the destaining solution, samples were incubated in 1 M KOH at 90 °C for 10 min, washed three times with distilled water, and incubated overnight in 1 ml of staining solution (10× phosphate-buffered saline: 1.4 M NaCl, 27 mM KCl, 10 mM Na₂HPO₄ × 2H₂O, and 18 mM KH₂PO₄, pH 7.3) with 0.2 µg/ml Alexa Fluor 488 dye (Thermo Fisher Scientific Inc., Waltham, MA, USA) in the dark. Samples were then washed three times with distilled water and examined using an Axioskop fluorescence microscope (Carl Zeiss AG, Oberkochen, Germany).

### Statistical analysis and visualisation of inoculation experiments

Statistical analyses and data visualisation were performed with R version 4.3.2 (R Core Team, 2023) using the following packages: dplyr (Wickham et al., 2023), tidyr (Wickham et al., 2024), ggplot2 (Wickham, 2016), stringr (Wickham, 2023), gridExtra (Auguie and Antonov, 2017), and inti (Lozano-Isla, 2025).

For the detached leaf assay, the number of necrotic spots per dm² was calculated for each leaf at 7, 10, and 14 dpi by dividing the total number of necrotic spots by the corresponding leaf area (dm²). To reflect steady or progressive disease development, the number of necrotic spots per dm² was not permitted to decrease over time. In cases where a lower value was recorded, the previous higher value was retained. This adjustment occurred when large necrotic areas developed, preventing further detection of individual spots by the YOLOv5s model. The percentage of necrotic leaf area was determined by subtracting the percentage of healthy leaf tissue, as assessed using Leaf_color_detector.py (Supplementary script S3), from 100. Subsequently, the mean of the four leaves per genotype-treatment-repetition combination was calculated for each time point. The area under the disease progress curve (AUDPC) was calculated using these mean values (Jeger and Viljanen-Rollinson, 2001). The AUDPC was calculated using the following formula:

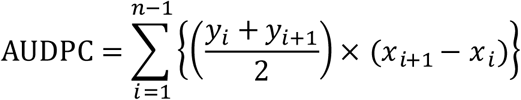

where *n* represents the total number of observations, *x* the time point of the observation, *y* the observed value, and *i* the index of each time point.

For the greenhouse experiment, the mean phenotyping score of the three leaves per genotype-treatment-repetition combination was calculated at each time point. These mean scores were then used for AUDPC calculation using the same formula as described above. Additionally, mean data were generated across repetitions per year and across all repetitions for both the detached leaf assay and the greenhouse experiment.

Spearman correlation coefficients were calculated to assess the relationship between individual repetitions and the two phenotyping methods. Broad-sense heritability was estimated using the robust Cullis method (Cullis et al., 2006; Covarrubias-Pazaran, 2019) via the H2cal function of the inti package (Lozano-Isla, 2025). Estimates were calculated for each trait and time point in the F_1_ population across all repetitions from 2023 and 2025, using the mean value of each genotype per repetition for both the detached leaf assay (n = 4) and the greenhouse experiment (n = 5).

### QTL analysis

QTL analysis was performed with MapQTL 5 (van Ooijen, 2004) using simple interval mapping. A permutation test with 1,000 iterations at a 95% confidence level was applied to determine the genome-wide significance threshold. For each linkage group (LG), only the position with the highest logarithm of the odds (LOD) score was considered, and QTLs were reported only if this maximum LOD score reached this threshold. In addition, a Kruskal-Wallis test was applied to averaged datasets that had shown significant results in simple interval mapping across all repetitions.

### Haplotype mapping of QTLs and KEGG pathway analysis of QTL regions

To assign the identified genome-wide significant QTLs across all repetitions to one of the two haplotypes of *Mb*j (Pfeifer et al., 2026b), the 122 F_1_ individuals were divided into two groups according to the alleles of the SNP markers with the highest LOD scores on the respective LGs. The presence of the QTL was attributed to the allele associated with lower apple blotch symptoms. Using the Basic Local Alignment Search Tool (Altschul et al., 1990) in CLC Main Workbench 25.0 (Qiagen, Venlo, The Netherlands), the SNP marker sequences were then aligned to the two haplotypes of *Mb*j, thereby allowing the haplotype carrying the resistance-associated allele, to be identified.

Functional characterisation of QTL regions was performed using Kyoto Encyclopedia of Genes and Genomes (KEGG) pathway analysis with KEGG Mapper (Kanehisa and Sato, 2020; Kanehisa et al., 2022). QTL regions were defined as the genomic intervals flanked by the nearest upstream and downstream SNP markers surrounding the marker with the highest LOD score, where the LOD value decreased by at least one unit relative to the maximum. Accordingly, the QTL regions spanned from HT1_LG01_23982724 to HT1_LG01_25975175 on LG 1, from HT1_LG02_8462643 to HT1_LG02_9246339 on LG 2, from HT1_LG12_18826669 to HT1_LG12_22021139 on LG 12, and from HT1_LG13_12131086 to HT1_LG13_15722331 on LG 13. KEGG annotation data for *Mb*j were obtained from Pfeifer et al. (2026b).

## Results

### Apple blotch resistance is polygenic and affected by phenotyping method and environment

In this study, a total of 466 datasets from the F_1_ population ’Idared’ × *Mb*j were used for simple interval mapping. Across all experiments and time points, 176 genome-wide significant associations were identified, distributed across 15 of the 17 LGs of *Mb*j. A summary overview is provided in Table 1, while all 176 significant associations detected in the detached leaf assay and greenhouse experiment are listed in Tables S2 and S3, respectively.

**Table 1.** Overview of the experimental design and simple interval mapping results of the F_1_ population.

| Phenotyping method | Trait | Description | Number of datasets | Number of significant associations | LGs with significant associations |
| --- | --- | --- | --- | --- | --- |
| Detached leaf assay | Necrotic spots per dm<sup>2</sup> | 3 time points (7, 10, 14 dpi) from 2 leaf origins (field and greenhouse) and mean of repetitions | 36 | 11 | 3, 5, 8, 9, 11, 15 |
|  | AUDPC of necrotic spots per dm<sup>2</sup> |  | 36 | 11 | 3, 5, 8, 9 |
|  | Percentage of necrotic leaf area |  | 36 | 3 | 16 |
|  | AUDPC of percentage necrotic leaf area |  | 36 | 4 | 10, 16 |
|  | Microscopy (five different characteristics) | 1 time point (21 dpi) from 2 leaf origins (field and greenhouse) | 10 | 1 | 14 |
| Greenhouse experiment | Phenotyping score | 12 time points (14-91 dpi) from 2 treatments (with and without zip bag) and mean of repetitions | 156 | 38 | 1, 2, 3, 4, 6, 12, 13, 14 |
|  | AUDPC of phenotyping score |  | 156 | 108 | 1, 2, 4, 13, 14 |
| Total across all experiments | / | / | 466 | 176 | 1, 2, 3, 4, 5, 6, 8, 9, 10, 11, 12, 13, 14, 15, 16 |

AUDPC: area under the disease progress curve; dpi: days post-inoculation; LG: linkage group.

From Table 1, it is evident that both phenotyping methods and all assessed traits yielded at least one significant QTL, indicating that each is suitable for evaluating different levels of apple blotch resistance. Using the detached leaf assay, QTLs were identified on LGs 3, 5, 8, 9, 11 and 15 for necrotic spots per dm²; on LGs 10 and 16 for percentage of necrotic leaf area; and on LG 14 based on microscopy. From the greenhouse experiment, QTLs on LGs 1, 2, 3, 4, 6, 12, 13, and 14 were detected based on the phenotyping score.

Only one QTL on LG 14 overlapped between the two experiments. This QTL was identified in the detached leaf assay based on the microscopic trait of the presence or absence of necrosis with acervuli surrounded by dark-brown tissue (see Figure 4d). The QTL was also detected at a similar position in the greenhouse experiment. All other microscopy-based mapping approaches did not lead to the detection of QTLs.

Traits based on the AUDPC resulted in a substantially higher number of significant associations compared with raw trait values (123 associations vs. 53). However, AUDPC-based traits were associated with QTLs distributed across a smaller number of LGs (11 vs. 14). Figures 5 and 6 show the inoculation results and Spearman correlation coefficients of the F_1_ population based on AUDPC values of the traits necrotic spots per dm^2^ and the phenotyping score at the final phenotyping time points of 14 and 91 dpi for the detached leaf assay and the greenhouse experiment, respectively. These AUDPC values represent the most comprehensive data, as they integrate disease development from the first to the last phenotyping time point and therefore provide an overall measure of disease progression.

**Figure 5.**
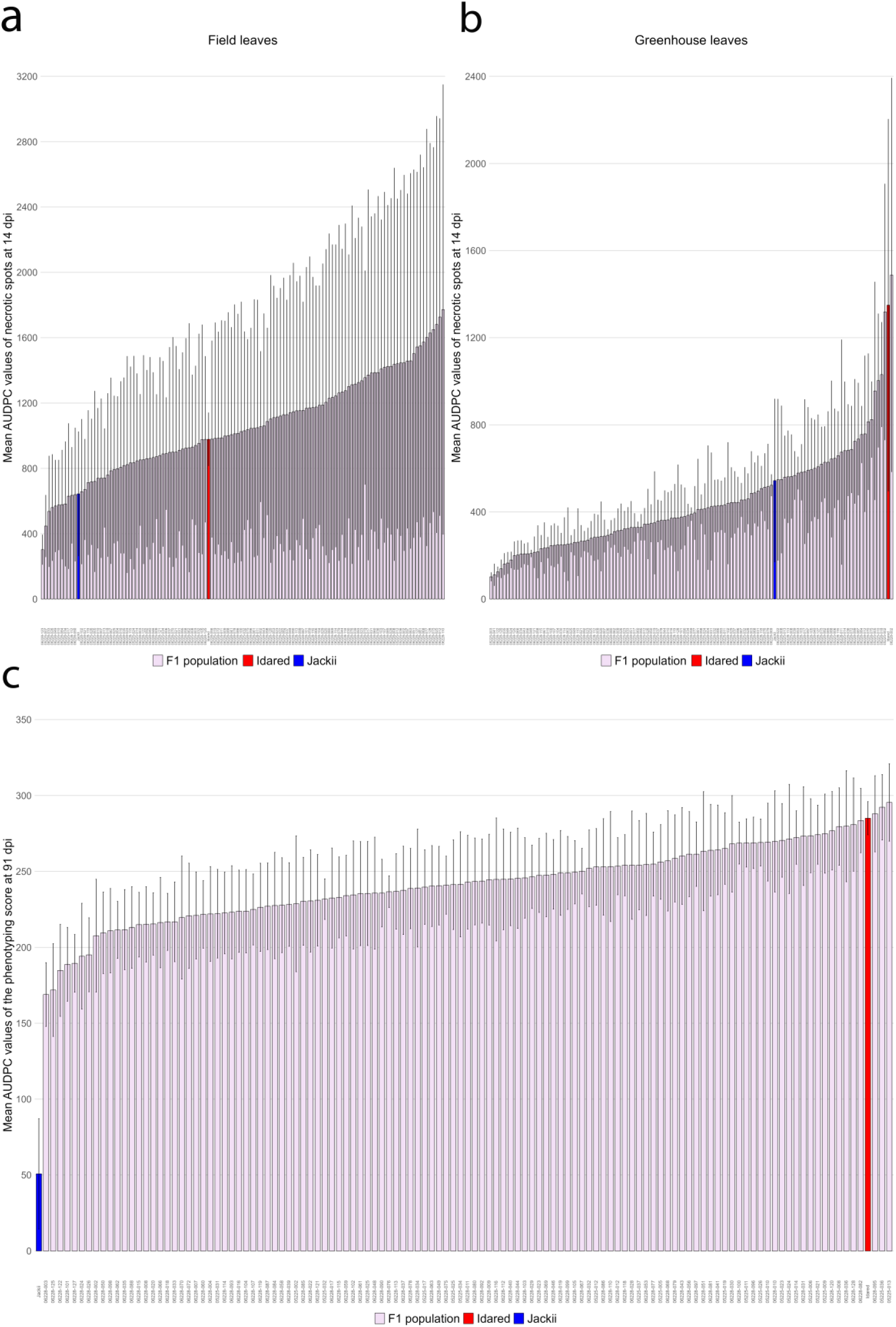
Inoculation results of the F_1_ population, including the parents ’Idared’ and *Mb*j. (a) Mean area under the disease progress curve (AUDPC) for the number of necrotic spots per dm² at 14 dpi in the detached leaf assay across all repetitions for field leaves; (b) the same for greenhouse leaves; (c) mean AUDPC values of the phenotyping score at 91 dpi in the greenhouse experiment across all repetitions with zip bag treatment. Error bars in all panels represent the standard error.

**Figure 6.**
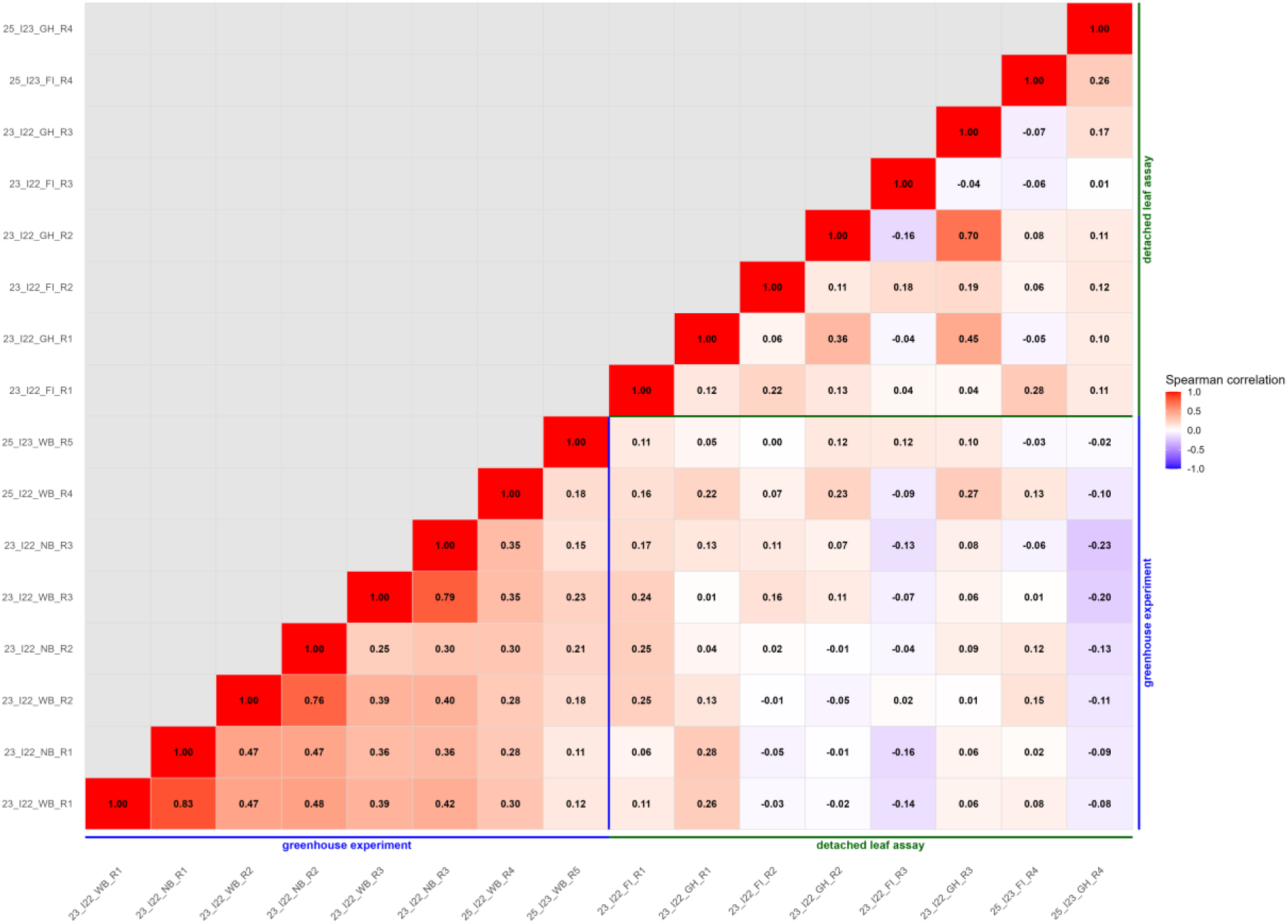
Spearman correlation coefficients of the F_1_ population derivedfrom the cross ’Idared’ × *Mb*j, calculated from the area under the disease progresscurve (AUDPC) at the final phenotyping time points for each repetition: number of necrotic spots per dm² at 14 dpi in the detachedleaf assay, and AUDPC of the phenotyping score at 91 dpi in the greenhouse experiment. Abbreviations: first two numbers = year of inoculation (22 = 2022, 23 = 2023); type of inoculum (I22 = inoculum from 2022, I23 = inoculum from 2023); treatment for greenhouse experiment (WB = with zip bag, NB = without zip bag) or origin of leaves for detached leaf assay (FI = field leaves, GH = greenhouse leaves); R = repetition number.

All F_1_ genotypes showed susceptibility to apple blotch in both phenotyping methods. Notably, the standard error was considerably higher in the detached leaf assay than in the greenhouse experiment and was higher for field leaves compared to greenhouse leaves. Ove rall, susceptibility showed substantial variability, which was, for example, evident for the parental genotypes in the detached leaf assay. Across all repetitions, the mean AUDPC of the F_1_ population in the detached leaf assay was 1061.97, with a standard error of 726.57 for field leaves, and 438.14 with a standard error of 154.82 for greenhouse leaves. In the greenhouse experiment, *Mb*j was consistently the most resistant genotype across all repetitions, showing no disease symptoms in the first three repetitions conducted in 2023 and an overall mean AUDPC of 50.75. By contrast, ’Idared’ was among the most susceptible genotypes, with a mean AUDPC of 285.02. With the exception of three genotypes, the F_1_ offspring showed susceptibility levels between the two parents, but generally closer to ’Idared’ than *Mb*j. The mean AUDPC of the F_1_ population across the five repetitions was 241.50, with a mean standard error of 27.37. Overall, the continuous distribution of susceptibility within the F_1_ population and the predominantly intermediate performance of the offspring between the two parental genotypes do not support the presence of a single major resistance locus, but rather indicate quantitative inheritance.

Spearman correlation coefficients between single repetitions of AUDPC values of necrotic spots per dm^2^ at 14 dpi from the detached leaf assay and phenotyping score at 91 dpi from the greenhouse experiment were generally below 0.50, with one exception in the detached leaf assay, where a correlation coefficient of 0.70 was observed. Overall, correlation coefficients within the greenhouse experiment were higher (mean 0.36) and exclusively positive compared with those from the detached leaf assay (mean 0.12). The three highest correlation coefficients (0.83, 0.79, and 0.76) were observed in the greenhouse experiment between the two treatments conducted with and without zip bags within the same repetition. These high correlations were consistent across all three repetitions in which both treatments were applied, indicating a robust and reproducible relationship. In contrast, the lowest correlations were observed between the two phenotyping methods, detached leaf assay and greenhouse experiment, with mean correlation coefficient of 0.05 and several negative values down to -0.23. Taken together, the generally low correlations between the two phenotyping methods indicate limited concordance in resistance assessment, whereas the high correlations between treatments within the same experimental repetition in the greenhouse experiment demonstrate a high level of inoculation consistency and reproducibility, irrespective of treatment.

In addition to Spearman correlation coefficients, Cullis broad-sense heritability was calculated for all traits assessed in the F_1_ population across all repetitions. In the detached leaf assay, Cullis broad-sense heritability ranged from 0.12 to 0.68, whereas in the greenhouse experiment it ranged from 0.18 to 0.60. For all traits and time points, residual variances consistently exceeded genetic variances. Detailed results are provided in Tables S4 and S5. The consistently higher residual relative to genetic variances further emphasises the substantial contribution of environmental and unexplained variance to the observed variation in resistance.

### Simple interval mapping with mean data across all repetitions reveals four genome-wide significant QTLs

To identify the most relevant QTLs, mean values across all four repetitions of the detached leaf assay and all five repetitions of the greenhouse experiment were calculated and used for simple interval mapping. QTLs that were significant at the genome-wide level across all repetitions are presented in Table 2 and Figure 7.

**Figure 7.**
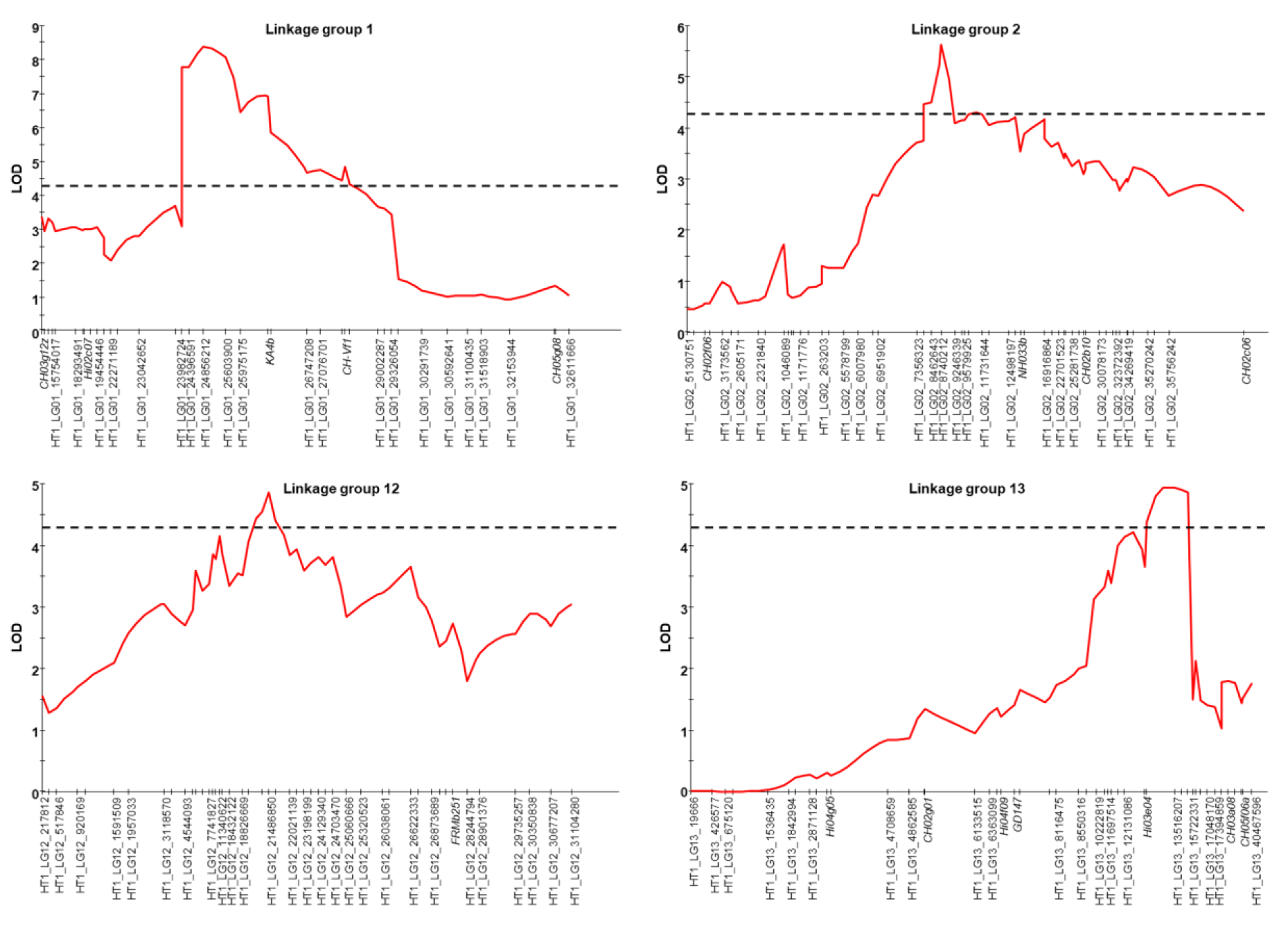
Highest logarithm of the odds (LOD) score profiles of LGs 1, 2, 12 and 13 identified by simple interval mapping using mean data across all five repetitions of the greenhouse experiment in the F_1_ population. The dashed lines indicate the genome-wide significance threshold determined by permutation testing with 1,000 iterations at a 95% confidence level.

**Table 2.**
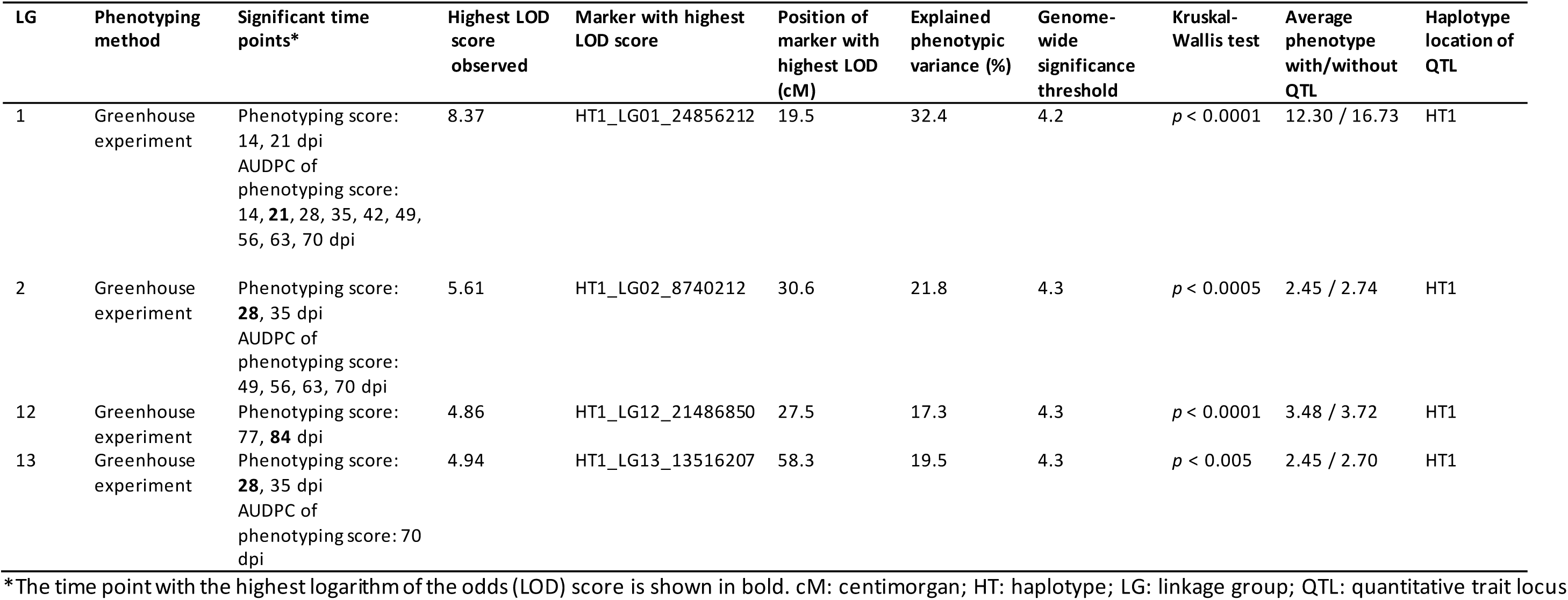
Genome-wide significant QTLs for apple blotch resistance detected by simple interval mapping using the mean across the 2023 and 2025 inoculation data for the F_1_ population.

| LG | Phenotyping method | Significant time points* | Highest LOD score observed | Marker with highest LOD score | Position of marker with highest LOD (cM) | Explained phenotypic variance (%) | Genome-wide significance threshold | Kruskal-Wallis test | Average phenotype with/without QTL | Haplotype location of QTL |
| --- | --- | --- | --- | --- | --- | --- | --- | --- | --- | --- |
| 1 | Greenhouse experiment | Phenotyping score: 14, 21 dpi<br>AUDPC of phenotyping score: 14, **21**, 28, 35, 42, 49, 56, 63, 70 dpi | 8.37 | HT1_LG01_24856212 | 19.5 | 32.4 | 4.2 | $p < 0.0001$ | 12.30 / 16.73 | HT1 |
| 2 | Greenhouse experiment | Phenotyping score: **28**, 35 dpi<br>AUDPC of phenotyping score: 49, 56, 63, 70 dpi | 5.61 | HT1_LG02_8740212 | 30.6 | 21.8 | 4.3 | $p < 0.0005$ | 2.45 / 2.74 | HT1 |
| 12 | Greenhouse experiment | Phenotyping score: 77, **84** dpi | 4.86 | HT1_LG12_21486850 | 27.5 | 17.3 | 4.3 | $p < 0.0001$ | 3.48 / 3.72 | HT1 |
| 13 | Greenhouse experiment | Phenotyping score: **28**, 35 dpi<br>AUDPC of phenotyping score: 70 dpi | 4.94 | HT1_LG13_13516207 | 58.3 | 19.5 | 4.3 | $p < 0.005$ | 2.45 / 2.70 | HT1 |

\*The time point with the highest logarithm of the odds (LOD) score is shown in bold. cM: centimorgan; HT: haplotype; LG: linkage group; QTL: quantitative trait locus

No genome-wide significant QTL was detected for the detached leaf assay when mean data across repetitions were analysed, whereas four QTLs were identified for the greenhouse experiment. Among these four QTLs, the strongest signal was detected on LG 1 based on AUDPC values derived from the phenotyping score at 21 dpi, with a LOD score of 8.37 and 32.4% phenotypic variance explained. The remaining three QTLs showed LOD scores and explained phenotypic variances of 5.61 and 21.8% on LG 2 at 28 dpi, 4.86 and 17.3% on LG 12 at 84 dpi, and 4.94 and 19.5% on LG 13 at 28 dpi, all based on the phenotyping score. In all cases, these four QTLs represented the highest LOD scores on their respective LG. In individual repetitions, LOD scores on these LGs were either comparable to those obtained from the mean data, as observed for LG 1, lower, as for LG 2, or not significant, as for LGs 12 and 13 (Table S3). Notably, all four QTLs were time-point specific. The highest LOD scores for QTLs on LGs 1, 2 and 13 were observed at 21-28 dpi, whereas the highest LOD score for LG 12 was detected at 84 dpi.

Haplotype analysis indicated that all QTLs are located on haplotype 1 of *Mb*j. KEGG pathway analysis revealed that genes associated with the mitogen-activated protein kinase (MAPK) signaling pathway (map04016), plant hormone signal transduction (map04075) and plant-pathogen interaction (map04626) were identified in each QTL region, highlighting their functional relevance in plant defence responses. Overall, robust genome-wide significant QTLs were identified only in the greenhouse experiment and were predominantly time-point specific, indicating that apple blotch resistance is both environment-dependent and dynamically regulated over the course of infection.

### Genotypes derived from open pollination exhibit lower resistance to apple blotch than *Mb*j

Following inoculation of the F_1_ population in the greenhouse experiment in 2023, all F_1_ descendants were found to be susceptible to apple blotch, whereas *Mb*j remained symptom-free. This observation suggested that resistance in *Mb*j may be conferred by a single recessive locus. To test this hypothesis, seeds from open-pollinated *Mb*j and two F_1_ descendants (05225-31 and 06228-66) were collected in autumn 2023. Genotyping with SSR markers revealed that the majority of these offspring had been pollinated by F_1_ individuals from the cross ’Idared’ × *Mb*j (Table 3). A subset of these genotypes was subsequently inoculated with *D. coronariae* in 2024.

**Table 3.** Overview of SSR genotyping results for open-pollinated genotypes.

| Population | Mother plant | Number of analysed genotypes | Identified outcrossers | Potential crossers with 'Idared' or *Mbj* | Observed self-fertilisations | Crossers with the F<sub>1</sub> population |
| --- | --- | --- | --- | --- | --- | --- |
| 'Jackii'-OA | *Mbj* | 232 | 22 | 2 | 0 | 208 |
| 05225-31-OA | 05225-31 | 40 | 0 | 2 | 0 | 38 |
| 06228-66-OA | 06228-66 | 30 | 1 | 5 | 0 | 24 |

In total, 20 genotypes from 05225-31 × F_1_, 15 genotypes of 06228-66 × F_1_, and 96 genotypes of *Mb*j × F_1_ were included in the detached leaf assay. For the greenhouse experiment, the same genotypes as in the detached leaf assay were used, with the addition of 65 further *Mb*j × F_1_ genotypes, bringing the total to 161. The results of these inoculations are shown in Figure 8.

**Figure 8.**
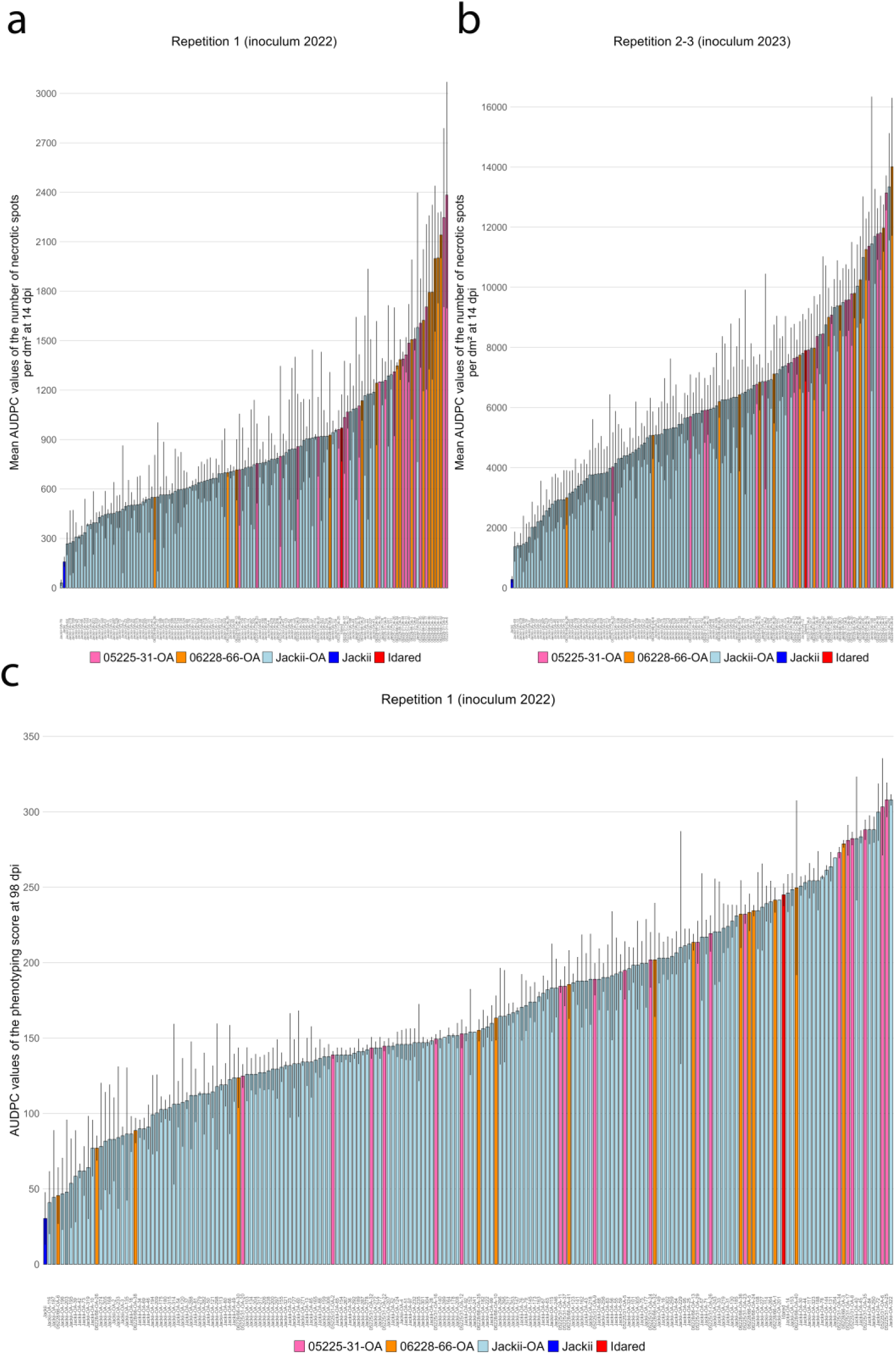
Inoculation results of open-pollinated genotypes, including ’Idared’ and *Mb*j, shown with standard error. (a) Area under the disease progress curve (AUDPC) for the number of necrotic spots per dm² at 14 dpi in the detached leaf assay for repetition 1 using inoculum from 2022; (b) the same based on the mean of repetitions 2 and 3 using inoculum from 2023; (c) AUDPC values of the phenotyping score at 98 dpi in the greenhouse experiment for repetition 1. If apple blotch resistance in *Mb*j were controlled by a single major recessive locus, approximately 50% of ‘Jackii-OA’ and 25% of 05225-31-OA and 06228-66-OA would be expected to display resistance levels comparable to *Mb*j; however, this pattern was not observed.

Genotypes derived from the F_1_ descendants 05225-31 and 06228-66 were generally more susceptible to apple blotch than those derived from *Mb*j in the detached leaf assay. Inoculations with inoculum from 2023 resulted in markedly higher symptom severity compared with inoculum from 2022, with mean AUDPC values of 858.14 and 6105.92, respectively, across the three populations. Nevertheless, the relative distribution of susceptibility among the three populations remained consistent across inoculum sources: 05225-31-OA and 06228-66-OA were consistently more susceptible than ‘Jackii’-OA.

In the greenhouse experiment, *Mb*j-derived genotypes were again less susceptible to apple blotch than offspring derived from the two F_1_ descendants. The ‘Jackii’-OA population exhibited a mean AUDPC value of 159.86 at 98 dpi, compared with 181.61 for 06228-66-OA and 210.47 for 05225-31-OA. *Mb*j remained the genotype with the lowest recorded symptoms in the greenhouse experiment, with an AUDPC value of 30.33, showing only slight symptoms. The segregation pattern across populations does not support the presence of a single major recessive resistance locus.

### Inoculum from 2023 differs from that of 2022 in virulence, genetic composition, and germination time

As previously noted, the inoculum from 2023 generally caused more disease severity than the inoculum from 2022. To further investigate these differences, the genetic composition of the inocula was determined. Only MLG 38 was detected in the 2022 inoculum, whereas both MLG 38 and MLG 23 were present in the inoculum from 2023. MLG 38 and MLG 23 differ by a single SSR marker out of 12. Germination rates of both inocula were consistently above 90% in all experiments; however, the inoculum from 2023 reached this rate after 24 h, whereas the inoculum from 2022 required 48 h, with almost no germination observed after 24 h, indicating a clear difference.

### Microscopic analyses show that *Mb*j is not immune to apple blotch

Microscopic examination of inoculated *Mb*j leaves showed characteristic infection structures of *D. coronariae*, including hyphae, subcuticular hyphal strands, haustoria, and acervuli with conidia (Figure 9). However, the quantity of disease symptoms was significantly lower than in susceptible genotypes, particularly in leaves from the greenhouse experiment. Conidia collected by scraping acervuli on *Mb*j, suspended in water, and then applied to ’Golden Delicious’ resulted in successful reinfection, demonstrating that the conidia produced on *Mb*j are both viable and virulent. These results demonstrate that *D. coronariae* can sporulate on *Mb*j after artificial inoculation, despite the overall reduced symptom severity.

**Figure 9.**
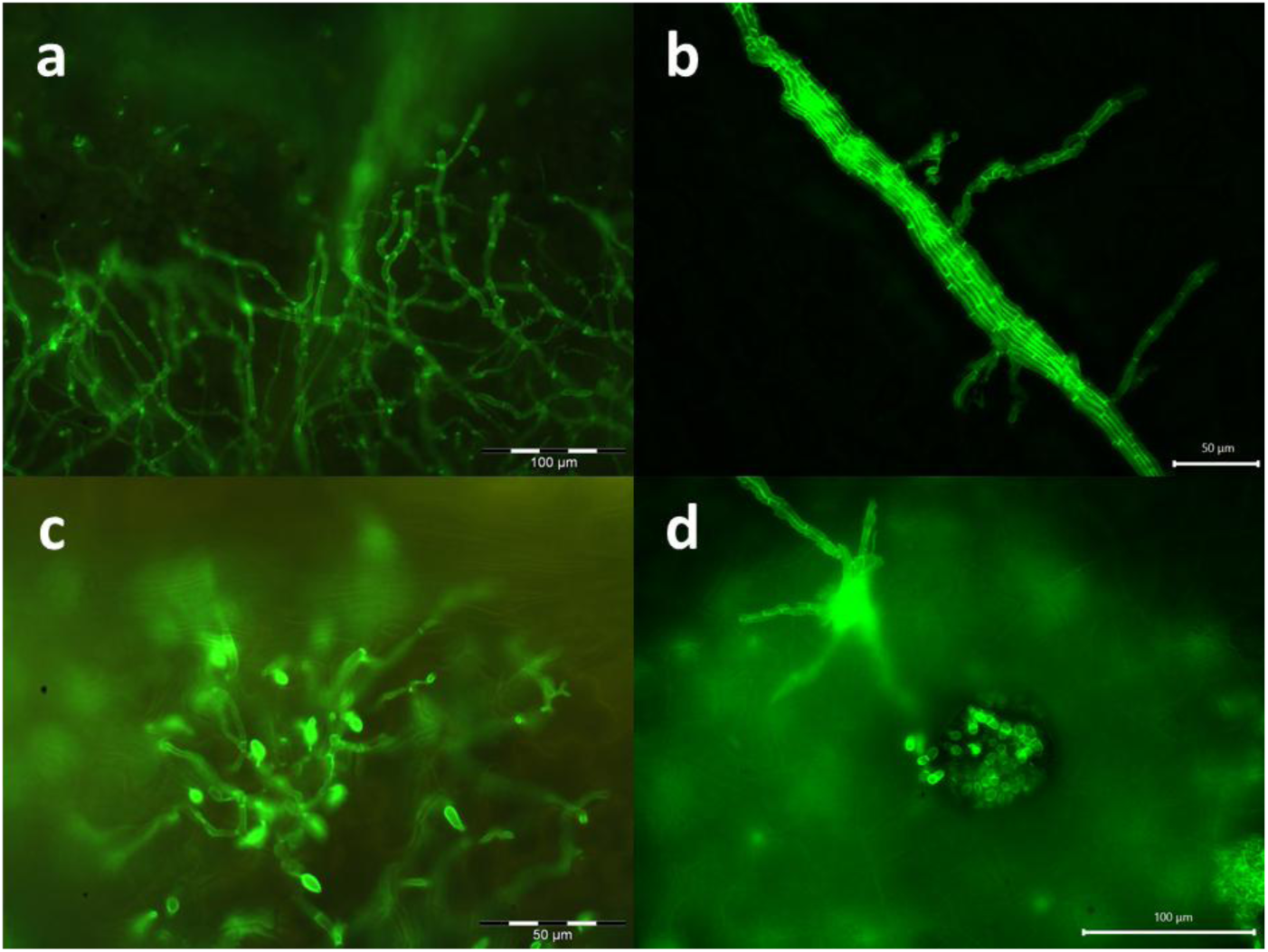
Microscopic analysis of *Mb*j leaves infected with *D. coronariae,* showing typical symptoms of susceptibility: (a) hyphae;(b)subcuticular hyphal strands;(c) hyphae with haustoria; (d)an acervulus with visible conidia.

## Discussion

The results of this study demonstrate that apple blotch resistance in *Mb*j is a complex quantitative trait influenced by multiple genomic loci. Genome-wide significant QTLs were detected on 15 out of 17 LGs; however, their detection was highly dependent on the phenotyping method, assessment time point, evaluated trait and environmental conditions. Together with the observation that none of the offspring derived from open pollination reached the resistance level of the resistance donor, these findings indicate that apple blotch resistance in *Mb*j is unlikely to be controlled by a single major dominant or recessive locus. Instead, this resistance most likely results from the combined action of multiple loci with small to moderate effects. Under the assumption of a single major recessive locus, approximately 25% of the F_1_ × F_1_ and 50% of the *Mb*j × F_1_ offspring would have been expected to display resistance levels comparable to *Mb*j.

A major outcome of this study is the strong influence of the phenotyping method on QTL detection. While both the detached leaf assay and the greenhouse experiment identified genome-wide significant QTLs, the detected LGs differed markedly between the two methods. Only LGs 3 and 14 were shared between the phenotyping methods, and among these, only the QTL on LG 14 showed a peak at a similar cM position. This strongly suggests that the two phenotyping methods capture different biological aspects of apple blotch resistance. The detached leaf assay likely quantifies localised responses, such as infection efficiency through necrotic spots and early lesion development, over a limited time span, followed by leaf senescence, whereas the greenhouse experiment integrates whole -plant effects and defence responses over a longer observation period. Although consistent results between inoculation assays of detached leaves and whole plants have been reported in other plant species (Abe et al., 2010; Miller-Butler et al., 2018), as well as in roses infected with *Diplocarpon rosae*, a fungus closely related to *D. coronariae* (Debener, 2019), a study in *Arabidopsis* demonstrated that, following inoculation with a hemibiotrophic pathogen, defence mechanisms can differ significantly between detached and attached leaves, with disease symptoms in detached leaves being associated with senescence (Liu et al., 2007). This may explain why our results differed markedly between the phenotyping methods, and similar findings have also been reported recently (Richter et al., 2025a).

AUDPC values, which integrate disease progression over time, resulted in QTL detection on fewer LGs compared to raw trait values and may have failed to capture some QTLs that are active only at specific infection stages. At the same time, AUDPC values revealed QTLs not detected using raw data, such as the QTL on LG 10 in the detached leaf assay, indicating that AUDPC can uncover biologically relevant loci that may otherwise remain undetected. Among all assessed traits, the number of necrotic spots per dm² in the detached leaf assay and the phenotyping score in the greenhouse experiment proved to be the most informative. In contrast, the percentage of necrotic leaf area in the detached leaf assay was considerably less informative, a finding that is consistent with previous studies (Richter et al., 2025a; Richter et al., 2025b). Moreover, this trait can be misleading, as necrotic leaf area is often triggered by secondary infections with other fungi. For this reason, detached leaves were phenotyped only up to 14 dpi, as non-specific tissue decay became evident at later time points. Although the highest Cullis broad-sense heritability value was observed for the trait percentage of necrotic leaf area at 7 dpi, this trait has limited biological relevance. The percentage of necrotic leaf area was generally very low at this early time point and, across all time points, resulted in only seven significant associations out of 176 detected overall. This underscores that heritability estimates should always be interpreted in conjunction with complementary analyses, as heritability alone often leads to misconceptions (Covarrubias-Pazaran, 2019).

Only four QTLs were identified using the mean data across all repetitions, and all of them originated from the greenhouse experiment. Notably, the LOD score of these QTLs was never exceeded in any single repetition and, in some cases, these QTLs were detected exclusively in the mean data. This suggests that the effects of these QTLs were stable, but their effects were masked by strong residual variation in individual repetitions. Increasing the number of biological replicates may therefore increase the power to detect such QTLs; in fire blight resistance phenotyping for example, up to ten biological replicates per genotype are commonly used (Peil et al., 2019; Emeriewen et al., 2021; Emeriewen et al., 2023), and in some studies even 20 or more have been used (Nybom et al., 2012; Piazza et al., 2021). As no stable QTLs were identified using mean values across all repetitions in the detached leaf assay, we conclude that the greenhouse experiment is more suitable for identifying apple blotch resistance loci. This conclusion is further supported by the higher variability observed in the detached leaf assay, reflected by lower mean correlation coefficients, relatively higher standard errors, and lower Cullis broad-sense heritability values, particularly when field-grown leaves were used, compared to the greenhouse experiment. This result was unexpected, as detached leaf assays are conducted under undoubtedly more consistent controlled conditions in Petri dishes than the greenhouse experiment. Notably, standard errors within the F_1_ population were substantially higher in detached leaf assays using field-grown leaves compared to greenhouse-grown leaves, although inoculation was performed in the same manner, on the same day, and with the same inoculum. The observation that the three highest correlation coefficients in the F_1_ population were found between the two treatments within the same repetition in the greenhouse experiment, while correlations between repetitions were substantially lower, suggests both that the inoculation procedure was accurate and reproducible and that the overall physiological status of the plant may have played an important role. Furthermore, the observation that relative susceptibility among genotypes for apple blotch resistance varied is not a new phenomenon and has also been reported in the literature, as certain genotypes were described as resistant in some studies but susceptible in others (Yin et al., 2013a; Khodadadi et al., 2022; Richter et al., 2025a). Taken together, these results indicate that both leaf- and plant-level physiological status contribute substantially to phenotypic variability to apple blotch resistance.

Previous studies have mapped apple blotch resistance primarily to chromosomes 9 and 12 (Noh et al., 2020; Richter et al., 2025b). Of the four QTLs identified in the present study using mean data, one is located on chromosome 12; however its genomic position differs from that reported previously (Richter et al., 2025b). This discrepancy is not surprising given that our analysis focused on *Mb*j rather than on a *M. domestica* collection. Within the four QTL regions, genes assigned to the KEGG pathways MAPK signaling, plant hormone signal transduction and plant-pathogen interaction were identified. Consistent with this, a previous study demonstrated that MAPK and Ca²⁺ signaling pathways were induced following inoculation with *D. coronariae*, and that the less susceptible *M. baccata* ’Shandingzi’ exhibited a higher proportion of disease resistance-related genes compared with ’Fuji’ (Feng et al., 2019). Other studies have highlighted the importance of the oxidative burst, salicylic acid signaling, chitinases, β-1,3-glucanases, and pathogenesis-related genes, such as *PR1* and *PR5*, which are typical defences against biotrophic pathogens, as well as jasmonic acid signaling at later stages of infection, which is typical of defence against necrotrophic pathogens, in resistance to the hemibiotrophic fungus *D. coronariae* (Yin et al., 2013b; Li et al., 2014; Sun et al., 2018; Liu et al., 2022; Khodadadi et al., 2022). In summary, the KEGG pathways identified align well with previously described defence mechanisms against *D. coronariae*. More recently, also effector proteins of *D. coronariae* have been investigated to gain insights into the molecular mechanisms by which the fungus manipulates its apple host (Guo et al., 2025) .

A correlation between resistance to apple blotch and *Alternaria alternata* has been observed, suggesting a shared genetic basis for resistance to these two diseases (Li et al., 2012a). We can partially confirm that resistance to other diseases may contribute to apple blotch resistance. In three of the 176 significant associations, the marker CH03e03 showed the highest LOD score. This marker is positioned at the top of chromosome 3, where resistance of *Mb*j to fire blight has also been localised and is associated with the *FB_Mr5* homolog, a gene encoding a CC-NBS-LRR resistance protein (Peil et al., 2007; Vogt et al., 2013; Fahrentrapp et al., 2013; Wöhner et al., 2014; Broggini et al., 2014; Wöhner et al., 2016) . However, it is more likely that another resistance gene at a nearby locus contributes to apple blotch resistance rather than the *FB_Mr5* homolog itself. Resistance genes are well known to occur in clusters in plant genomes (Young, 2000; Perazzolli et al., 2014).

Genetic analyses of the inocula from 2022 and 2023 revealed the presence of an additional MLG in the inoculum from 2023, which could potentially explain the observed differences in virulence between the inocula. However, this MLG differs at only 1 of 12 SSR markers (Dc68; MLG 38 has a fragment length of 352 bp, whereas MLG 23 has 334 bp), and the MLGs observed in Europe are described as genetically homogeneous and clonal (Oberhänsli et al., 2021). Furthermore, to date, no sexual reproduction of *D. coronariae* has been described in Europe, likely due to the potential absence of the MAT1-1 mating-type locus (Richter et al., 2024). We therefore consider it unlikely that the observed differences in virulence were primarily caused by MLG 23; however, variability in virulence among isolates within the same MLG cannot be entirely excluded and it has been described that long terminal repeat retrotransposons can contribute to genomic variation in *D. coronariae* (Gao et al., 2026) . Instead, the delayed germination of the inoculum from 2022 suggests a strong year effect on the inoculum, independent of the presence of different MLGs. Such year effects may be related to environmental factors, like the hot and dry summer of 2022 (Herrera-Lormendez et al., 2023), or to differences in harvest time. Although the use of cultured single-spore isolates under standardised conditions, along with improved cultivation media (Chauhan and Modgil, 2024) could potentially reduce this source of variability, cultivation of *D. coronariae* on artificial media negatively affects virulence (Wöhner et al., 2019). Therefore, this approach was deemed unsuitable for the present study. Nevertheless, the use of cultured and genetically transformed *D. coronariae* could be valuable in future analyses to better understand the interactions between the fungus and the apple host (Chauhan et al., 2021).

Microscopic analyses clearly demonstrated that *Mb*j can be successfully infected with *D. coronariae*. However, the observation of variable phenotypes, including the absence of visible symptoms in some repetitions, suggests that certain QTLs in *Mb*j may be active only under specific environmental conditions or at particular time points, indicating that different genetic mechanisms may be involved during early infection stages, and later disease development. This could also explain why no symptoms on this genotype were identified in previous studies (Wöhner et al., 2021; Pfeifer et al., 2024). Temporal or environmental specificity of QTLs associated with disease resistance has been reported in apple (Calenge and Durel, 2006; Karlström et al., 2022; Emeriewen et al., 2024) and in other plant species (Welz et al., 1999; Li et al., 2007; Li et al., 2012b). Currently, there is no evidence suggesting that resistance of *Mb*j to apple blotch is related to the cuticle, as abrasion did not lead to a higher detected amount of fungal DNA (Pfeifer et al., 2024). Although abrasion of cuticle has been reported to increase susceptibility to *D. rosae*, this effect was not observed in all genotypes (Castledine et al., 1981). It is also well established that artificial inoculations under controlled conditions often overestimate susceptibility and do not always align with field observations (Franke and Brenneman, 2001; Hampson and Sholberg, 2008; Delgado et al., 2022). This is likely also the case for apple blotch, as the environment had a strong influence on disease symptom development, with residual variance consistently exceeding genetic variance. Consistent with this, no apple blotch infection was observed in 2025 in the F_1_ population at the experimental orchard of the Julius Kühn-Institut (JKI), even though *M. domestica* trees in the neighboring rows were severely infected. These observations highlight the importance of field trials, despite their inherent disadvantages, such as higher costs, the need for multi-year experiments, uneven pathogen presence, the potential for misidentification of symptoms with those caused by other diseases, lower experimental control and possibly reduced reproducibility. Field inoculations with multiple replications may help mitigate some of these limitations. Nevertheless, SNPs associated with resistance to apple blotch were successfully identified using data from natural field infections, indicating that this is a suitable approach (Noh et al., 2020).

The findings of this study emphasise the difficulty of improving apple blotch resistance through conventional breeding, as the trait is polygenic in *Mb*j, and *Mb*j represents a wild genotype carrying numerous undesirable horticultural characteristics that would require extensive breeding efforts to eliminate (Flachowsky et al., 2011). While transgenic approaches have demonstrated that apple blotch resistance can be enhanced under laboratory conditions (Sun et al., 2018; Tan et al., 2020; Kumar et al., 2025), the identification of inherently resistant genotypes carrying a single major dominant locus would greatly facilitate breeding and therefore remains of high importance, particularly given that many consumers prefer conventionally bred apples over genetically modified ones. As most, if not all, *M. domestica* cultivars appear to be susceptible to apple blotch (Richter et al., 2025a), screening wild *Malus* accessions represents a valuable strategy for identifying novel sources of tolerance or resistance (Wöhner et al., 2021). In this context, the accessions MAL0778 and MAL0278 have so far shown no evidence of *D. coronariae* infection despite repeated inoculation experiments (data not shown) and should be further investigated in future studies. Until a single major resistance locus for apple blotch is identified, conventional breeding will likely require the pyramiding of multiple small-effect loci with cumulative effects.

## Conclusion

In this study, populations derived from the apple blotch-resistant genotype *Mb*j were artificially inoculated with *D. coronariae*. Segregation patterns in the F_1_ population and in populations derived from open pollination indicate that resistance in *Mb*j is not controlled by a single major locus. Instead, our results demonstrate a polygenic mode of inheritance, with QTLs identified on 15 of the 17 LGs. Most QTLs were detected only in a subset of the datasets, and susceptibility to *D. coronariae* was strongly influenced by inoculum, phenotyping method and environmental conditions. QTLs were identified at different time points, and those detected in the greenhouse experiment differed markedly from those in the detached leaf assay. QTLs detected using mean data across all repetitions were exclusively identified in the greenhouse experiment and were located on LGs 1, 2, 12 and 13, with the markers showing the highest LOD scores explaining between 17.3 and 32.4% of the phenotypic variance. Before substantial breeding efforts are undertaken, the effects of these QTLs should be validated under field conditions. In parallel, the search for genotypes carrying a single major resistance locus should be continued, as this would greatly facilitate the breeding of apple blotch-resistant cultivars.

## Acknowledgments

We gratefully thank Dr. Thomas Oberhänsli for providing primers for genotyping *D. coronariae*, Dr. Anne Bohr for providing *D. coronariae*-infected leaves, Sabine Bartsch and Jessica Göhler for technical assistance with inoculations, and the personnel responsible for plant care.

## AI statement

ChatGPT (OpenAI, https://openai.com) was used for language editing.

## Author contributions

MP: strategy and design, plant material, analyses and writing, review and editing. AP: plant material, review and editing. HF: conception, funding acquisition, strategy and design, supervision, review and editing. OFE: conception, review and editing. TWW: conception, funding acquisition, strategy and design, supervision, review and editing.

