## Supplementary Materials for "Inoculation of *Malus baccata* ‘Jackii’-derived offspring and QTL analysis reveal a polygenic inheritance pattern of apple blotch resistance"

### Supplementary Tables

**Table S1.** List of SSR markers employed for genotyping genotypes derived from open pollination

| Marker | Reference |
|---|---|
| CH01f02, CH01f03b, CH01f07a, CH01h01, CH02c09, CH02d08, CH02g09, CH03d07, CH05e03, CH05f06 | Liebhard et al., 2002 |
| GD 12 | Hokanson et al., 1998 |

**Table S2.** Genome-wide significant associations identified by simple interval mapping in the detached leaf assay across all datasets

| Trait | Year of inoculation | Inoculum | Leaf origin | Repetition | dpi | LG | Highest LOD score | Marker with highest LOD score | Position (cM) | Explained phenotypic variance (%) |
|---|---|---|---|---|---|---|---|---|---|---|
| **Necrotic spots per $dm^2$** | **2023** | **2022** | **Field** | **1** | **7** | **8** | **7.37** | **HT1_LG08_25126438** | **51.2** | **24.3** |
| Necrotic spots per $dm^2$ | 2023 | 2022 | Field | 1 | 10 | 8 | 5.46 | HT1_LG08_28047111 | 54.6 | 19.2 |
| **Necrotic spots per $dm^2$** | **2023** | **2022** | **Field** | **1** | **14** | **9** | **4.49** | **HT1_LG09_6055205** | **33.0** | **16.4** |
| Necrotic spots per $dm^2$ | 2023 | 2022 | Greenhouse | 2 | 10 | 3 | 3.30 | CH03e03 | 0.0 | 12.0 |
| **Necrotic spots per $dm^2$** | **2023** | **2022** | **Greenhouse** | **2** | **14** | **3** | **4.86** | **CH03e03** | **0.0** | **17.2** |
| **Necrotic spots per $dm^2$** | **2023** | **2022** | **Greenhouse** | **2** | **14** | **11** | **4.00** | **HT1_LG11_34294231** | **45.2** | **20.2** |
| **Necrotic spots per $dm^2$** | **2023** | **2022** | **Greenhouse** | **1-3** | **7** | **5** | **4.95** | **HT1_LG05_24210086** | **19.0** | **21.5** |
| Necrotic spots per $dm^2$ | 2023 | 2022 | Greenhouse | 1-3 | 10 | 5 | 4.06 | HT1_LG05_24210086 | 19.0 | 15.6 |
| Necrotic spots per $dm^2$ | 2023 | 2022 | Greenhouse | 1-3 | 10 | 11 | 4.05 | HT1_LG11_34294231 | 45.2 | 20.9 |
| **Necrotic spots per $dm^2$** | **2023** | **2022** | **Greenhouse** | **1-3** | **14** | **11** | **4.28** | **HT1_LG11_34294231** | **45.2** | **21.3** |
| **Necrotic spots per $dm^2$** | **2025** | **2023** | **Field** | **4** | **14** | **15** | **4.93** | **HT1_LG15_41074458** | **79.2** | **17.4** |
| **AUDPC of necrotic spots per $dm^2$** | **2023** | **2022** | **Field** | **1** | **7** | **8** | **7.37** | **HT1_LG08_25126438** | **51.2** | **24.3** |
| AUDPC of necrotic spots per $dm^2$ | 2023 | 2022 | Field | 1 | 10 | 8 | 7.02 | HT1_LG08_25126438 | 51.2 | 23.4 |
| AUDPC of necrotic spots per $dm^2$ | 2023 | 2022 | Field | 1 | 10 | 9 | 4.40 | HT1_LG09_5217202 | 32.0 | 16.3 |
| AUDPC of necrotic spots per $dm^2$ | 2023 | 2022 | Field | 1 | 14 | 8 | 5.53 | HT1_LG08_28047111 | 54.6 | 19.3 |
| **AUDPC of necrotic spots per $dm^2$** | **2023** | **2022** | **Field** | **1** | **14** | **9** | **4.45** | **HT1_LG09_6055205** | **33.0** | **16.5** |
| **AUDPC of necrotic spots per $dm^2$** | **2023** | **2022** | **Field** | **1-3** | **10** | **8** | **4.70** | **HT1_LG08_25126438** | **51.2** | **16.3** |
| **AUDPC of necrotic spots per $dm^2$** | **2023** | **2022** | **Greenhouse** | **2** | **10** | **5** | **4.16** | **HT1_LG05_24210086** | **19.0** | **16.5** |
| **AUDPC of necrotic spots per $dm^2$** | **2023** | **2022** | **Greenhouse** | **2** | **14** | **3** | **3.82** | **CH03e03** | **0.0** | **13.8** |
| **AUDPC of necrotic spots per $dm^2$** | **2023** | **2022** | **Greenhouse** | **3** | **10** | **5** | **5.00** | **HT1_LG05_6790436** | **3.7** | **17.8** |
| AUDPC of necrotic spots per $dm^2$ | 2023 | 2022 | Greenhouse | 1-3 | 7 | 5 | 4.95 | HT1_LG05_24210086 | 19.0 | 21.5 |
| **AUDPC of necrotic spots per $dm^2$** | **2023** | **2022** | **Greenhouse** | **1-3** | **10** | **5** | **6.04** | **HT1_LG05_24210086** | **19.0** | **25.0** |
| Percentage of necrotic leaf area | 2023 | 2022 | Greenhouse | 3 | 7 | 16 | 5.00 | HT1_LG16_16651738 | 57.5 | 23.8 |
| **Percentage of necrotic leaf area** | **2023** | **2022** | **Greenhouse** | **3** | **10** | **16** | **5.04** | **HT1_LG16_16651738** | **57.5** | **23.9** |
| Percentage of necrotic leaf area | 2023 | 2022 | Greenhouse | 3 | 14 | 16 | 4.61 | HT1_LG16_16651738 | 57.5 | 21.9 |
| **AUDPC of percentage of necrotic leaf area** | **2023** | **2022** | **Field** | **1-3** | **14** | **10** | **4.78** | **FRM4** | **27.2** | **18.0** |
| AUDPC of percentage of necrotic leaf area | 2023 | 2022 | Greenhouse | 3 | 7 | 16 | 5.00 | HT1_LG16_16651738 | 57.5 | 23.8 |
| AUDPC of percentage of necrotic leaf area | 2023 | 2022 | Greenhouse | 3 | 10 | 16 | 5.02 | HT1_LG16_16651738 | 57.5 | 23.9 |
| **AUDPC of percentage of necrotic leaf area** | **2023** | **2022** | **Greenhouse** | **3** | **14** | **16** | **5.11** | **HT1_LG16_16651738** | **57.5** | **24.2** |

| | | | | | | | | | | |
|---|---|---|---|---|---|---|---|---|---|---|
| **Presence/absence of necrosis with acervuli surrounded by dark-brown tissue** | **2023** | **2022** | **Field** | **3** | **21** | **14** | **5.90** | **HT1_LG14_28796794** | **54.0** | **23.8** |

For each unique combination, the row corresponding to the marker with the highest logarithm of the odds (LOD) score per linkage group (LG) across all time points is highlighted in bold. AUDPC: area under the disease progress curve; cM: centimorgan; dpi: days post-inoculation.

**Table S3.** Genome-wide significant associations identified by simple interval mapping in the greenhouse experiment across all datasets

| Trait | Year of inoculation | Inoculum | Treatment | Repetition | dpi | LG | Highest LOD score | Marker with highest LOD score* | Position (cM) | Explained phenotypic variance (%) |
|---|---|---|---|---|---|---|---|---|---|---|
| Phenotyping score | 2023 | 2022 | With zip bag | 1 | 28 | 14 | 4.39 | HT1_LG14_19111015 | 19.3 | 16.7 |
| Phenotyping score | 2023 | 2022 | With zip bag | 1 | 35 | 14 | 4.52 | HT1_LG14_27543259 | 49.2 | 16.4 |
| **Phenotyping score** | **2023** | **2022** | **With zip bag** | **1** | **42** | **14** | **4.74** | **HT1_LG14_27543259** | **49.2** | **17.2** |
| **Phenotyping score** | **2023** | **2022** | **With zip bag** | **2** | **14** | **1** | **7.66** | **HT1_LG01_24114628** | **16.9** | **37.7** |
| Phenotyping score | 2023 | 2022 | With zip bag | 2 | 21 | 1 | 4.89 | HT1_LG01_24856212 | 19.5 | 20.9 |
| **Phenotyping score** | **2023** | **2022** | **With zip bag** | **3** | **21** | **1** | **4.38** | **HT1_LG01_29924205** | **43** | **73.5** |
| **Phenotyping score** | **2023** | **2022** | **With zip bag** | **1-3** | **14** | **1** | **7.67** | **HT1_LG01_24396591** | **17.8** | **38.7** |
| Phenotyping score | 2023 | 2022 | With zip bag | 1-3 | 21 | 1 | 6.22 | HT1_LG01_24856212 | 19.5 | 27.9 |
| Phenotyping score | 2023 | 2022 | With zip bag | 1-3 | 28 | 1 | 4.53 | HT1_LG01_29002287 | 40.5 | 21.8 |
| **Phenotyping score** | **2023** | **2022** | **With zip bag** | **1-3** | **28** | **3** | **4.50** | **HT1_LG03_37007502** | **78.6** | **19.0** |
| **Phenotyping score** | **2023** | **2022** | **With zip bag** | **1-3** | **35** | **14** | **4.95** | **HT1_LG14_27543259** | **49.2** | **17.6** |
| Phenotyping score | 2023 | 2022 | With zip bag | 1-3 | 42 | 14 | 4.80 | HT1_LG14_27543259 | 49.2 | 17.1 |
| **Phenotyping score** | **2023** | **2022** | **With zip bag** | **1-3** | **70** | **4** | **5.04** | **Hi07b02** | **30.0** | **18.9** |
| **Phenotyping score** | **2023** | **2022** | **Without zip bag** | **1** | **21** | **14** | **4.53** | **HT1_LG14_19111015** | **19.3** | **17.5** |
| Phenotyping score | 2023 | 2022 | Without zip bag | 2 | 35 | 2 | 5.18 | HT1_LG02_8740212 | 30.6 | 20.4 |
| **Phenotyping score** | **2023** | **2022** | **Without zip bag** | **2** | **42** | **2** | **5.39** | **HT1_LG02_8740212** | **30.6** | **21.4** |
| **Phenotyping score** | **2023** | **2022** | **Without zip bag** | **3** | **28** | **2** | **4.47** | **HT1_LG02_16066858** | **43.0** | **20.4** |
| Phenotyping score | 2023 | 2022 | Without zip bag | 1-3 | 21 | 14 | 4.42 | HT1_LG14_28024105 | 52.3 | 15.9 |
| Phenotyping score | 2023 | 2022 | Without zip bag | 1-3 | 28 | 14 | 4.54 | HT1_LG14_29010646 | 55.8 | 17.7 |
| Phenotyping score | 2023 | 2022 | Without zip bag | 1-3 | 35 | 1 | 4.29 | HT1_LG01_25603900 | 22.2 | 17.0 |
| **Phenotyping score** | **2023** | **2022** | **Without zip bag** | **1-3** | **35** | **14** | **5.09** | **HT1_LG14_28024105 (4.93)** | **52.3** | **19.8** |
| **Phenotyping score** | **2023** | **2022** | **Without zip bag** | **1-3** | **42** | **1** | **4.38** | **HT1_LG01_25603900** | **22.2** | **17.9** |
| **Phenotyping score** | **2025** | **2022** | **With zip bag** | **4** | **77** | **6** | **4.33** | **HT1_LG06_11098342 (4.23)** | **14.3** | **16.4** |
| Phenotyping score | 2025 | 2022, 2023 | With zip bag | 4-5 | 14 | 4 | 4.63 | HT1_LG04_7914653 | 13.6 | 18.0 |
| **Phenotyping score** | **2025** | **2022, 2023** | **With zip bag** | **4-5** | **35** | **4** | **4.73** | **HT1_LG04_23637976** | **27.1** | **18.3** |
| **Phenotyping score** | **2025** | **2022, 2023** | **With zip bag** | **4-5** | **35** | **13** | **4.65** | **HT1_LG13_14598871** | **60.3** | **19.7** |
| **Phenotyping score** | **2023, 2025** | **2022** | **With zip bag** | **1-4** | **14** | **1** | **6.89** | **HT1_LG01_24396591** | **17.8** | **32.0** |
| Phenotyping score | 2023, 2025 | 2022 | With zip bag | 1-4 | 21 | 1 | 4.94 | HT1_LG01_25603900 (4.93) | 20.5 | 21.5 |
| **Phenotyping score** | **2023, 2025** | **2022** | **With zip bag** | **1-4** | **28** | **13** | **4.33** | **HT1_LG13_14598871** | **60.3** | **16.7** |
| Phenotyping score | 2023, 2025 | 2022 | With zip bag | 1-4 | 35 | 13 | 4.20 | HT1_LG13_13516207 | 58.7 | 16.2 |

| | | | | | | | | | | |
|---|---|---|---|---|---|---|---|---|---|---|
| **Phenotyping score** | **2023, 2025** | **2022, 2023** | **With zip bag** | **1-5** | **14** | **1** | **7.95** | **HT1_LG01_24856212** | **19.5** | **30.8** |
| Phenotyping score | 2023, 2025 | 2022, 2023 | With zip bag | 1-5 | 21 | 1 | 6.14 | HT1_LG01_25603900 (6.12) | 21.5 | 24.0 |
| **Phenotyping score** | **2023, 2025** | **2022, 2023** | **With zip bag** | **1-5** | **28** | **2** | **5.61** | **HT1_LG02_8740212** | **30.6** | **21.8** |
| **Phenotyping score** | **2023, 2025** | **2022, 2023** | **With zip bag** | **1-5** | **28** | **13** | **4.94** | **HT1_LG13_13516207 (4.93)** | **58.3** | **19.5** |
| Phenotyping score | 2023, 2025 | 2022, 2023 | With zip bag | 1-5 | 35 | 2 | 4.50 | HT1_LG02_8740212 | 30.6 | 17.7 |
| Phenotyping score | 2023, 2025 | 2022, 2023 | With zip bag | 1-5 | 35 | 13 | 4.36 | HT1_LG13_14253854 | 59.4 | 18.0 |
| Phenotyping score | 2023, 2025 | 2022, 2023 | With zip bag | 1-5 | 77 | 12 | 4.52 | HT1_LG12_21486850 | 27.5 | 16.2 |
| **Phenotyping score** | **2023, 2025** | **2022, 2023** | **With zip bag** | **1-5** | **84** | **12** | **4.86** | **HT1_LG12_21486850** | **27.5** | **17.3** |
| AUDPC of phenotyping score | 2023 | 2022 | With zip bag | 1 | 28 | 14 | 4.26 | HT1_LG14_19111015 | 19.3 | 16.2 |
| AUDPC of phenotyping score | 2023 | 2022 | With zip bag | 1 | 35 | 14 | 4.55 | HT1_LG14_19111015 | 19.3 | 17.2 |
| AUDPC of phenotyping score | 2023 | 2022 | With zip bag | 1 | 42 | 14 | 4.41 | HT1_LG14_19111015 | 19.3 | 16.7 |
| AUDPC of phenotyping score | 2023 | 2022 | With zip bag | 1 | 49 | 14 | 4.52 | HT1_LG14_27543259 | 49.2 | 16.4 |
| AUDPC of phenotyping score | 2023 | 2022 | With zip bag | 1 | 56 | 14 | 4.65 | HT1_LG14_19111015 | 19.3 | 17.6 |
| **AUDPC of phenotyping score** | **2023** | **2022** | **With zip bag** | **1** | **63** | **14** | **4.85** | **HT1_LG14_19111015** | **19.3** | **18.3** |
| AUDPC of phenotyping score | 2023 | 2022 | With zip bag | 1 | 70 | 14 | 4.84 | HT1_LG14_19111015 | 19.3 | 18.3 |
| AUDPC of phenotyping score | 2023 | 2022 | With zip bag | 1 | 77 | 14 | 4.69 | HT1_LG14_19111015 | 19.3 | 17.8 |
| AUDPC of phenotyping score | 2023 | 2022 | With zip bag | 1 | 84 | 14 | 4.5 | HT1_LG14_19111015 | 19.3 | 17.1 |
| AUDPC of phenotyping score | 2023 | 2022 | With zip bag | 1 | 91 | 14 | 4.32 | HT1_LG14_19111015 | 19.3 | 16.5 |
| AUDPC of phenotyping score | 2023 | 2022 | With zip bag | 2 | 14 | 1 | 7.66 | HT1_LG01_24114628 | 16.9 | 37.7 |
| **AUDPC of phenotyping score** | **2023** | **2022** | **With zip bag** | **2** | **21** | **1** | **8.37** | **HT1_LG01_24396591** | **17.8** | **39.1** |
| AUDPC of phenotyping score | 2023 | 2022 | With zip bag | 2 | 28 | 1 | 7.89 | HT1_LG01_24396591 | 17.8 | 37.1 |
| AUDPC of phenotyping score | 2023 | 2022 | With zip bag | 2 | 35 | 1 | 6.96 | HT1_LG01_24396591 | 17.8 | 32.7 |
| AUDPC of phenotyping score | 2023 | 2022 | With zip bag | 2 | 42 | 1 | 6.11 | HT1_LG01_24396591 | 17.8 | 28.3 |
| AUDPC of phenotyping score | 2023 | 2022 | With zip bag | 2 | 49 | 1 | 5.69 | HT1_LG01_25603900 | 22.2 | 22.7 |
| AUDPC of phenotyping score | 2023 | 2022 | With zip bag | 2 | 49 | 14 | 4.21 | HT1_LG14_28024105 | 52.3 | 15.5 |
| AUDPC of phenotyping score | 2023 | 2022 | With zip bag | 2 | 56 | 1 | 5.50 | HT1_LG01_25603900 | 22.2 | 21.6 |
| AUDPC of phenotyping score | 2023 | 2022 | With zip bag | 2 | 63 | 1 | 5.31 | HT1_LG01_25603900 | 22.2 | 20.8 |
| AUDPC of phenotyping score | 2023 | 2022 | With zip bag | 2 | 63 | 14 | 4.27 | HT1_LG14_27543259 | 49.2 | 15.7 |

| | | | | | | | | | | |
|---|---|---|---|---|---|---|---|---|---|---|
| AUDPC of phenotyping score | 2023 | 2022 | With zip bag | 2 | 70 | 1 | 5.13 | HT1_LG01_25603900 | 22.2 | 20.1 |
| AUDPC of phenotyping score | 2023 | 2022 | With zip bag | 2 | 77 | 1 | 5.00 | HT1_LG01_25603900 | 22.2 | 19.5 |
| AUDPC of phenotyping score | 2023 | 2022 | With zip bag | 2 | 84 | 1 | 4.92 | HT1_LG01_25603900 | 22.2 | 19.2 |
| AUDPC of phenotyping score | 2023 | 2022 | With zip bag | 2 | 91 | 1 | 4.86 | HT1_LG01_25603900 | 22.2 | 19.0 |
| **AUDPC of phenotyping score** | **2023** | **2022** | **With zip bag** | **2** | **91** | **14** | **4.30** | **HT1_LG14_27543259** | **49.2** | **15.8** |
| AUDPC of phenotyping score | 2023 | 2022 | With zip bag | 1-3 | 14 | 1 | 7.67 | HT1_LG01_24396591 | 17.8 | 38.7 |
| **AUDPC of phenotyping score** | **2023** | **2022** | **With zip bag** | **1-3** | **21** | **1** | **8.25** | **HT1_LG01_24396591** | **17.8** | **40.7** |
| AUDPC of phenotyping score | 2023 | 2022 | With zip bag | 1-3 | 28 | 1 | 8.16 | HT1_LG01_24396591 | 17.8 | 40.3 |
| AUDPC of phenotyping score | 2023 | 2022 | With zip bag | 1-3 | 35 | 1 | 7.76 | HT1_LG01_24396591 | 17.8 | 38.5 |
| AUDPC of phenotyping score | 2023 | 2022 | With zip bag | 1-3 | 35 | 14 | 4.38 | HT1_LG14_28024105 | 52.3 | 15.7 |
| AUDPC of phenotyping score | 2023 | 2022 | With zip bag | 1-3 | 35 | 14 | 4.31 | HT1_LG14_27543259 | 49.2 | 15.5 |
| AUDPC of phenotyping score | 2023 | 2022 | With zip bag | 1-3 | 42 | 1 | 7.28 | HT1_LG01_24396591 | 17.8 | 36.1 |
| AUDPC of phenotyping score | 2023 | 2022 | With zip bag | 1-3 | 42 | 14 | 4.83 | HT1_LG14_27543259 | 49.2 | 17.2 |
| AUDPC of phenotyping score | 2023 | 2022 | With zip bag | 1-3 | 49 | 1 | 6.91 | HT1_LG01_25603900 (6.90) | 21.5 | 30.06 |
| AUDPC of phenotyping score | 2023 | 2022 | With zip bag | 1-3 | 49 | 14 | 5.15 | HT1_LG14_27543259 | 49.2 | 18.2 |
| AUDPC of phenotyping score | 2023 | 2022 | With zip bag | 1-3 | 56 | 1 | 6.72 | HT1_LG01_25603900 | 22.2 | 27.8 |
| AUDPC of phenotyping score | 2023 | 2022 | With zip bag | 1-3 | 56 | 14 | 5.37 | HT1_LG14_27543259 | 49.2 | 18.9 |
| AUDPC of phenotyping score | 2023 | 2022 | With zip bag | 1-3 | 63 | 1 | 6.55 | HT1_LG01_25603900 | 22.2 | 27.0 |
| AUDPC of phenotyping score | 2023 | 2022 | With zip bag | 1-3 | 63 | 14 | 5.52 | HT1_LG14_27543259 | 49.2 | 19.4 |
| AUDPC of phenotyping score | 2023 | 2022 | With zip bag | 1-3 | 70 | 1 | 6.38 | HT1_LG01_25603900 | 22.2 | 26.1 |
| AUDPC of phenotyping score | 2023 | 2022 | With zip bag | 1-3 | 70 | 14 | 5.60 | HT1_LG14_27543259 | 49.2 | 19.6 |
| AUDPC of phenotyping score | 2023 | 2022 | With zip bag | 1-3 | 77 | 1 | 6.24 | HT1_LG01_25603900 | 22.2 | 25.5 |
| AUDPC of phenotyping score | 2023 | 2022 | With zip bag | 1-3 | 77 | 14 | 5.65 | HT1_LG14_27543259 | 49.2 | 19.8 |
| AUDPC of phenotyping score | 2023 | 2022 | With zip bag | 1-3 | 84 | 1 | 6.12 | HT1_LG01_25603900 | 22.2 | 24.9 |
| **AUDPC of phenotyping score** | **2023** | **2022** | **With zip bag** | **1-3** | **84** | **14** | **5.66** | **HT1_LG14_27543259** | **49.2** | **19.8** |
| AUDPC of phenotyping score | 2023 | 2022 | With zip bag | 1-3 | 91 | 1 | 6.00 | HT1_LG01_25603900 | 22.2 | 24.4 |

| | | | | | | | | | | |
|---|---|---|---|---|---|---|---|---|---|---|
| AUDPC of phenotyping score | 2023 | 2022 | With zip bag | 1-3 | 91 | 14 | 5.63 | HT1_LG14_27543259 | 49.2 | 19.7 |
| AUDPC of phenotyping score | 2023 | 2022 | Without zip bag | 1 | 49 | 14 | 4.33 | HT1_LG14_27543259 | 49.2 | 15.5 |
| AUDPC of phenotyping score | 2023 | 2022 | Without zip bag | 1 | 56 | 14 | 4.49 | HT1_LG14_27543259 | 49.2 | 16.1 |
| **AUDPC of phenotyping score** | **2023** | **2022** | **Without zip bag** | **1** | **63** | **14** | **4.57** | **HT1_LG14_27543259** | **49.2** | **16.3** |
| AUDPC of phenotyping score | 2023 | 2022 | Without zip bag | 1 | 70 | 14 | 4.54 | HT1_LG14_27543259 | 49.2 | 16.2 |
| AUDPC of phenotyping score | 2023 | 2022 | Without zip bag | 1 | 77 | 14 | 4.44 | HT1_LG14_27543259 | 49.2 | 15.9 |
| AUDPC of phenotyping score | 2023 | 2022 | Without zip bag | 1 | 84 | 14 | 4.36 | HT1_LG14_27543259 | 49.2 | 15.7 |
| AUDPC of phenotyping score | 2023 | 2022 | Without zip bag | 1 | 91 | 14 | 4.35 | HT1_LG14_27543259 | 49.2 | 15.6 |
| AUDPC of phenotyping score | 2023 | 2022 | Without zip bag | 2 | 42 | 2 | 4.83 | HT1_LG02_8740212 (4.73) | 30.4 | 18.7 |
| **AUDPC of phenotyping score** | **2023** | **2022** | **Without zip bag** | **2** | **49** | **2** | **5.15** | **HT1_LG02_8740212 (5.10)** | **30.4** | **19.8** |
| AUDPC of phenotyping score | 2023 | 2022 | Without zip bag | 2 | 56 | 2 | 5.02 | HT1_LG02_8740212 | 30.6 | 19.5 |
| AUDPC of phenotyping score | 2023 | 2022 | Without zip bag | 2 | 63 | 2 | 4.91 | HT1_LG02_8740212 | 30.6 | 19.0 |
| AUDPC of phenotyping score | 2023 | 2022 | Without zip bag | 2 | 70 | 2 | 4.77 | HT1_LG02_8740212 | 30.6 | 18.5 |
| AUDPC of phenotyping score | 2023 | 2022 | Without zip bag | 2 | 77 | 2 | 4.57 | HT1_LG02_8740212 | 30.6 | 17.7 |
| AUDPC of phenotyping score | 2023 | 2022 | Without zip bag | 2 | 84 | 2 | 4.38 | HT1_LG02_8740212 | 30.6 | 17.0 |
| AUDPC of phenotyping score | 2023 | 2022 | Without zip bag | 2 | 91 | 2 | 4.21 | HT1_LG02_8893221 | 30.6 | 16.4 |
| AUDPC of phenotyping score | 2023 | 2022 | Without zip bag | 1-3 | 42 | 1 | 4.48 | HT1_LG01_24856212 | 19.5 | 19.9 |
| AUDPC of phenotyping score | 2023 | 2022 | Without zip bag | 1-3 | 42 | 14 | 4.80 | HT1_LG14_28024105 | 52.3 | 17.1 |
| **AUDPC of phenotyping score** | **2023** | **2022** | **Without zip bag** | **1-3** | **49** | **1** | **4.57** | **HT1_LG01_24856212** | **19.5** | **20.1** |
| AUDPC of phenotyping score | 2023 | 2022 | Without zip bag | 1-3 | 49 | 14 | 4.97 | HT1_LG14_28024105 | 52.3 | 17.6 |
| AUDPC of phenotyping score | 2023 | 2022 | Without zip bag | 1-3 | 56 | 1 | 4.52 | HT1_LG01_24856212 | 19.5 | 19.8 |
| AUDPC of phenotyping score | 2023 | 2022 | Without zip bag | 1-3 | 56 | 14 | 5.06 | HT1_LG14_28024105 | 52.3 | 17.9 |
| AUDPC of phenotyping score | 2023 | 2022 | Without zip bag | 1-3 | 63 | 1 | 4.46 | HT1_LG01_24856212 | 19.5 | 19.5 |
| **AUDPC of phenotyping score** | **2023** | **2022** | **Without zip bag** | **1-3** | **63** | **14** | **5.09** | **HT1_LG14_28024105** | **52.3** | **18.0** |
| AUDPC of phenotyping score | 2023 | 2022 | Without zip bag | 1-3 | 70 | 1 | 4.41 | HT1_LG01_24856212 | 19.5 | 19.2 |
| AUDPC of phenotyping score | 2023 | 2022 | Without zip bag | 1-3 | 70 | 14 | 5.06 | HT1_LG14_28024105 | 52.3 | 17.9 |

| | | | | | | | | | | |
|---|---|---|---|---|---|---|---|---|---|---|
| AUDPC of phenotyping score | 2023 | 2022 | Without zip bag | 1-3 | 77 | 14 | 5.02 | HT1_LG14_28024105 | 52.3 | 17.8 |
| AUDPC of phenotyping score | 2023 | 2022 | Without zip bag | 1-3 | 84 | 1 | 4.34 | HT1_LG01_24856212 | 19.5 | 18.7 |
| AUDPC of phenotyping score | 2023 | 2022 | Without zip bag | 1-3 | 84 | 14 | 4.97 | HT1_LG14_28024105 | 52.3 | 17.6 |
| AUDPC of phenotyping score | 2023 | 2022 | Without zip bag | 1-3 | 91 | 1 | 4.30 | HT1_LG01_24856212 | 19.5 | 18.4 |
| AUDPC of phenotyping score | 2023 | 2022 | Without zip bag | 1-3 | 91 | 14 | 4.90 | HT1_LG14_28024105 | 52.3 | 17.4 |
| **AUDPC of phenotyping score** | **2025** | **2022, 2023** | **With zip bag** | **4-5** | **14** | **4** | **4.63** | **HT1_LG04_7914653** | **13.6** | **18.0** |
| AUDPC of phenotyping score | 2023, 2025 | 2022 | With zip bag | 1-4 | 14 | 1 | 6.89 | HT1_LG01_24396591 | 17.8 | 32.0 |
| **AUDPC of phenotyping score** | **2023, 2025** | **2022** | **With zip bag** | **1-4** | **21** | **1** | **6.98** | **HT1_LG01_24856212** | **19.5** | **30.0** |
| AUDPC of phenotyping score | 2023, 2025 | 2022 | With zip bag | 1-4 | 28 | 1 | 6.51 | HT1_LG01_24856212 | 19.5 | 28.0 |
| AUDPC of phenotyping score | 2023, 2025 | 2022 | With zip bag | 1-4 | 35 | 1 | 5.85 | HT1_LG01_24856212 (5.82) | 20.5 | 25.1 |
| AUDPC of phenotyping score | 2023, 2025 | 2022 | With zip bag | 1-4 | 42 | 1 | 5.19 | HT1_LG01_25603900 (5.18) | 21.5 | 21.9 |
| AUDPC of phenotyping score | 2023, 2025 | 2022 | With zip bag | 1-4 | 42 | 13 | 4.32 | HT1_LG13_13516207 | 58.7 | 16.7 |
| AUDPC of phenotyping score | 2023, 2025 | 2022 | With zip bag | 1-4 | 42 | 14 | 4.2 | HT1_LG14_27543259 | 49.2 | 15.1 |
| AUDPC of phenotyping score | 2023, 2025 | 2022 | With zip bag | 1-4 | 49 | 1 | 4.53 | HT1_LG01_25603900 (4.53) | 21.5 | 19.3 |
| AUDPC of phenotyping score | 2023, 2025 | 2022 | With zip bag | 1-4 | 49 | 13 | 4.45 | HT1_LG13_13516207 | 58.7 | 17.2 |
| AUDPC of phenotyping score | 2023, 2025 | 2022 | With zip bag | 1-4 | 49 | 14 | 4.38 | HT1_LG14_27543259 | 49.2 | 15.7 |
| **AUDPC of phenotyping score** | **2023, 2025** | **2022** | **With zip bag** | **1-4** | **56** | **13** | **4.46** | **HT1_LG13_13516207** | **58.7** | **17.2** |
| **AUDPC of phenotyping score** | **2023, 2025** | **2022** | **With zip bag** | **1-4** | **56** | **14** | **4.42** | **HT1_LG14_27543259** | **49.2** | **15.8** |
| **AUDPC of phenotyping score** | **2023, 2025** | **2022** | **With zip bag** | **1-4** | **63** | **13** | **4.46** | **HT1_LG13_13516207** | **58.7** | **17.1** |
| AUDPC of phenotyping score | 2023, 2025 | 2022 | With zip bag | 1-4 | 63 | 14 | 4.33 | HT1_LG14_27543259 | 49.2 | 15.6 |
| AUDPC of phenotyping score | 2023, 2025 | 2022 | With zip bag | 1-4 | 70 | 13 | 4.39 | HT1_LG13_13516207 | 58.7 | 16.9 |
| AUDPC of phenotyping score | 2023, 2025 | 2022 | With zip bag | 1-4 | 77 | 13 | 4.20 | HT1_LG13_13516207 | 58.7 | 16.2 |
| AUDPC of phenotyping score | 2023, 2025 | 2022, 2023 | With zip bag | 1-5 | 14 | 1 | 7.95 | HT1_LG01_24856212 | 19.5 | 30.8 |
| **AUDPC of phenotyping score** | **2023, 2025** | **2022, 2023** | **With zip bag** | **1-5** | **21** | **1** | **8.37** | **HT1_LG01_24856212** | **19.5** | **32.4** |
| AUDPC of phenotyping score | 2023, 2025 | 2022, 2023 | With zip bag | 1-5 | 28 | 1 | 7.80 | HT1_LG01_24856212 (7.77) | 20.5 | 30.4 |
| AUDPC of phenotyping score | 2023, 2025 | 2022, 2023 | With zip bag | 1-5 | 35 | 1 | 6.80 | HT1_LG01_25603900 (6.76) | 21.5 | 26.2 |
| AUDPC of phenotyping score | 2023, 2025 | 2022, 2023 | With zip bag | 1-5 | 42 | 1 | 5.97 | HT1_LG01_25603900 (5.96) | 21.5 | 23.2 |
| AUDPC of phenotyping score | 2023, 2025 | 2022, 2023 | With zip bag | 1-5 | 49 | 1 | 5.38 | HT1_LG01_25603900 | 22.2 | 20.6 |
| AUDPC of phenotyping score | 2023, 2025 | 2022, 2023 | With zip bag | 1-5 | 49 | 2 | 4.32 | HT1_LG02_8740212 | 30.6 | 16.6 |

| | | | | | | | | | | |
|---|---|---|---|---|---|---|---|---|---|---|
| AUDPC of phenotyping score | 2023, 2025 | 2022, 2023 | With zip bag | 1-5 | 56 | 1 | 4.86 | HT1_LG01_25603900 | 22.2 | 18.8 |
| **AUDPC of phenotyping score** | **2023, 2025** | **2022, 2023** | **With zip bag** | **1-5** | **56** | **2** | **4.40** | **HT1_LG02_8740212** | **30.6** | **17.0** |
| AUDPC of phenotyping score | 2023, 2025 | 2022, 2023 | With zip bag | 1-5 | 63 | 1 | 4.44 | HT1_LG01_25603900 | 22.2 | 17.3 |
| AUDPC of phenotyping score | 2023, 2025 | 2022, 2023 | With zip bag | 1-5 | 63 | 2 | 4.39 | HT1_LG02_8740212 | 30.6 | 17.1 |
| AUDPC of phenotyping score | 2023, 2025 | 2022, 2023 | With zip bag | 1-5 | 70 | 1 | 4.13 | HT1_LG01_25603900 | 22.2 | 16.2 |
| AUDPC of phenotyping score | 2023, 2025 | 2022, 2023 | With zip bag | 1-5 | 70 | 2 | 4.27 | HT1_LG02_8740212 | 30.6 | 16.8 |
| **AUDPC of phenotyping score** | **2023, 2025** | **2022, 2023** | **With zip bag** | **1-5** | **70** | **13** | **4.11** | **HT1_LG13_14253854** | **59.4** | **16.7** |

For each unique combination, the row corresponding to the marker with the highest logarithm of the odds (LOD) score per linkage group (LG) across all time points is highlighted in bold. * LOD score of the respective marker is given in parentheses if a higher interpolated LOD score exists. AUDPC: area under the disease progress curve; cM: centimorgan; dpi: days post-inoculation.

**Table S4.** Cullis broad-sense heritability estimates for the $F_1$ population across all repetitions of the detached leaf assay

| Trait | Leaf origin | dpi | Cullis broad-sense heritability | Genetic variance | Residual variance |
|---|---|---|---|---|---|
| Necrotic spots per $dm^2$ | Field | 7 | 0.1504041 | 41.75649 | 928.4264 |
| Necrotic spots per $dm^2$ | Field | 10 | 0.1191345 | 238.1744 | 6934.096 |
| Necrotic spots per $dm^2$ | Field | 14 | 0.1198467 | 243.0972 | 7029.588 |
| Necrotic spots per $dm^2$ | Greenhouse | 7 | 0.4336041 | 50.48661 | 258.1869 |
| Necrotic spots per $dm^2$ | Greenhouse | 10 | 0.4460849 | 402.5932 | 1956.298 |
| Necrotic spots per $dm^2$ | Greenhouse | 14 | 0.4638069 | 950.0199 | 4296.528 |
| AUDPC of necrotic spots per $dm^2$ | Field | 7 | 0.1504041 | 511.517 | 11373.22 |
| AUDPC of necrotic spots per $dm^2$ | Field | 10 | 0.1498459 | 2762.225 | 61683.53 |
| AUDPC of necrotic spots per $dm^2$ | Field | 14 | 0.1335724 | 11199.76 | 286001.7 |
| AUDPC of necrotic spots per $dm^2$ | Greenhouse | 7 | 0.4336041 | 618.461 | 3162.789 |
| AUDPC of necrotic spots per $dm^2$ | Greenhouse | 10 | 0.4449213 | 3715.189 | 18138.15 |
| AUDPC of necrotic spots per $dm^2$ | Greenhouse | 14 | 0.4507839 | 23598.64 | 112497.7 |
| Percentage of necrotic leaf area | Field | 7 | 0.6771201 | 0.2453238 | 0.4526842 |
| Percentage of necrotic leaf area | Field | 10 | 0.2875233 | 0.2150126 | 2.092825 |
| Percentage of necrotic leaf area | Field | 14 | 0.1353804 | 2.272969 | 57.16085 |
| Percentage of necrotic leaf area | Greenhouse | 7 | 0.6585808 | 0.9076465 | 1.822341 |
| Percentage of necrotic leaf area | Greenhouse | 10 | 0.662712 | 1.029062 | 2.027813 |
| Percentage of necrotic leaf area | Greenhouse | 14 | 0.5385138 | 1.62573 | 5.433614 |
| AUDPC of percentage necrotic leaf area | Field | 7 | 0.6771201 | 3.005216 | 5.545382 |

| | | | | | |
|---|---|---|---|---|---|
| AUDPC of percentage necrotic leaf area | Field | 10 | 0.62585 | 10.16564 | 23.60406 |
| AUDPC of percentage necrotic leaf area | Field | 14 | 0.2035001 | 26.69398 | 411.2589 |
| AUDPC of percentage necrotic leaf area | Greenhouse | 7 | 0.6585808 | 11.11867 | 22.32367 |
| AUDPC of percentage necrotic leaf area | Greenhouse | 10 | 0.6602171 | 39.4135 | 78.54931 |
| AUDPC of percentage necrotic leaf area | Greenhouse | 14 | 0.6480424 | 114.7325 | 241.525 |

AUDPC: area under the disease progress curve; dpi: days post-inoculation.

**Table S5.** Cullis broad-sense heritability estimates for the $F_1$ population across all repetitions of the greenhouse experiment

| Trait | dpi | Cullis broad-sense heritability | Genetic variance | Residual variance |
|---|---|---|---|---|
| Phenotyping score | 14 | 0.374682 | 0.072363986 | 0.5674629 |
| Phenotyping score | 21 | 0.5811445 | 0.18111184 | 0.6071584 |
| Phenotyping score | 28 | 0.4847523 | 0.09942568 | 0.4942194 |
| Phenotyping score | 35 | 0.4702627 | 0.08540369 | 0.450214 |
| Phenotyping score | 42 | 0.4236039 | 0.06060667 | 0.3867077 |
| Phenotyping score | 49 | 0.3222033 | 0.03526517 | 0.3491716 |
| Phenotyping score | 56 | 0.3434378 | 0.0311252 | 0.2798583 |
| Phenotyping score | 63 | 0.320749 | 0.02522438 | 0.2514346 |
| Phenotyping score | 70 | 0.3158693 | 0.02263535 | 0.2307946 |
| Phenotyping score | 77 | 0.2819356 | 0.01813437 | 0.2176707 |
| Phenotyping score | 84 | 0.2639052 | 0.01579507 | 0.2077453 |
| Phenotyping score | 91 | 0.1751597 | 0.00788327 | 0.1754817 |
| AUDPC of phenotyping score | 14 | 0.374682 | 3.545835 | 27.80568 |
| AUDPC of phenotyping score | 21 | 0.4977997 | 19.440615 | 91.66156 |
| AUDPC of phenotyping score | 28 | 0.5713502 | 49.320402 | 172.20314 |
| AUDPC of phenotyping score | 35 | 0.590868 | 82.783943 | 266.40992 |
| AUDPC of phenotyping score | 42 | 0.5951207 | 119.077998 | 376.39379 |
| AUDPC of phenotyping score | 49 | 0.5865858 | 151.852811 | 497.52425 |
| AUDPC of phenotyping score | 56 | 0.5740541 | 179.946997 | 621.20886 |
| AUDPC of phenotyping score | 63 | 0.5734046 | 210.772316 | 729.57462 |
| AUDPC of phenotyping score | 70 | 0.5797834 | 245.624423 | 827.92343 |
| AUDPC of phenotyping score | 77 | 0.5852426 | 281.084204 | 926.06177 |
| AUDPC of phenotyping score | 84 | 0.5903743 | 317.043305 | 1022.26974 |
| AUDPC of phenotyping score | 91 | 0.5934786 | 350.406133 | 1115.16393 |

AUDPC: area under the disease progress curve; dpi: days post-inoculation.

**Supplementary scripts**

**Supplementary script S1. Leaf_area_calculator.py**

```
""" @author: Matthias Pfeifer """


import cv2
import numpy as np
import os
import glob
import pandas as pd
from concurrent.futures import ThreadPoolExecutor

# Function to resize the image
def resize_image(image, scale_percent=40):
    width = int(image.shape[1] * scale_percent / 100)
    height = int(image.shape[0] * scale_percent / 100)
    return cv2.resize(image, (width, height), interpolation=cv2.INTER_AREA)

# Function to extract the leaf from the image
def extract_leaf(image):
    lower_green = np.array([30, 50, 50])
    upper_green = np.array([90, 255, 255])
    lower_brown = np.array([0, 20, 7])
    upper_brown = np.array([40, 94, 30])
    lower_leaf = np.array([min(lower_green[0], lower_brown[0]),
                           min(lower_green[1], lower_brown[1]),
                           min(lower_green[2], lower_brown[2])])
    upper_leaf = np.array([max(upper_green[0], upper_brown[0]),
                           max(upper_green[1], upper_brown[1]),
                           max(upper_green[2], upper_brown[2])])

    hsv = cv2.cvtColor(image, cv2.COLOR_BGR2HSV)
    mask = cv2.inRange(hsv, lower_leaf, upper_leaf)
    contours, _ = cv2.findContours(mask, cv2.RETR_EXTERNAL, cv2.CHAIN_APPROX_SIMPLE)

    if contours:
        max_contour = max(contours, key=cv2.contourArea)
        cv2.drawContours(image, [max_contour], -1, (0, 255, 0), thickness=10)
        return image, max_contour
    else:
        print("No contours found for the image.")
        return None, None

# Function to find a rectangular structure in the image (adjusted for light blue)
def find_rectangular_structure(image):
    # Convert to HSV
    hsv = cv2.cvtColor(image, cv2.COLOR_BGR2HSV)

    # Define the color range for light blue (adjust if needed)
    lower_blue = np.array([80, 50, 50])
    upper_blue = np.array([130, 255, 255])
```

```
    # Create a mask for the blue color
    mask = cv2.inRange(hsv, lower_blue, upper_blue)

    # Find contours in the masked image
    contours, _ = cv2.findContours(mask, cv2.RETR_EXTERNAL, cv2.CHAIN_APPROX_SIMPLE)

    tolerance = 0.1  # 10% Tolerance for approximation
    best_rect = None
    best_area = 0

    for contour in contours:
        epsilon = tolerance * cv2.arcLength(contour, True)
        approx = cv2.approxPolyDP(contour, epsilon, True)

        if len(approx) == 4:
            x, y, w, h = cv2.boundingRect(approx)
            aspect_ratio = w / float(h)

            # Check if the contour is roughly rectangular
            if 0.9 <= aspect_ratio <= 1.1:
                area = w * h
                total_area = image.shape[0] * image.shape[1]

                # Ensure the rectangle is neither too small nor too large
                if 0.01 * total_area <= area <= 0.30 * total_area and area > best_area:
                    best_area = area
                    best_rect = (x, y, w, h)

    if best_rect is not None:
        x, y, w, h = best_rect
        cv2.rectangle(image, (x, y), (x + w, y + h), (0, 0, 255), 10)
        return image, best_rect
    else:
        print("No rectangular structure found.")
        return None, None

input_folder = 'input_images'
output_folder = 'output_images_leafarea'

def process_image(image_path):
    image = cv2.imread(image_path)
    image = resize_image(image, scale_percent=40)
    piece_image, piece_bbox = find_rectangular_structure(image)
    leaf_image, leaf_contour = extract_leaf(image)

    filename = os.path.basename(image_path)
    output_filename = os.path.splitext(filename)[0] + '_piece.jpg'
    output_path = os.path.join(output_folder, output_filename)
    cv2.imwrite(output_path, image, [int(cv2.IMWRITE_JPEG_QUALITY), 40])  # Set JPG quality to 40

    print(output_filename + " saved.")

    return filename, leaf_contour, piece_bbox
```

```
os.makedirs(output_folder, exist_ok=True)

# Limit the number of threads to 10
max_threads = 10
with ThreadPoolExecutor(max_workers=max_threads) as executor:
    processed_images = list(executor.map(process_image, glob.glob(os.path.join(input_folder, '*.jpg'))))

print("Process completed.")

data = {'Image_name': [], 'Piecearea': [], 'Leafarea': [], 'Leafarea_calculated': []}

for filename, leaf_contour, piece_bbox in processed_images:
    if leaf_contour is not None and piece_bbox is not None:
        data['Image_name'].append(filename)

        leaf_area = cv2.contourArea(leaf_contour)
        x, y, w, h = piece_bbox
        piece_area = w * h
        data['Piecearea'].append(piece_area)
        data['Leafarea'].append(leaf_area)
        data['Leafarea_calculated'].append((leaf_area / piece_area) * 4.00)
    else:
        data['Image_name'].append(filename)
        data['Piecearea'].append("NA")
        data['Leafarea'].append("NA")
        data['Leafarea_calculated'].append("NA")

df = pd.DataFrame(data)
excel_file = 'Output-file.xlsx'
df.to_excel(excel_file, index=False)
print("Excel file created", excel_file
```

**Supplementary script 2. Leaf_extractor_resizer.py**

```
""" @author: Matthias Pfeifer """



import cv2
import numpy as np
import os
import glob
import openpyxl
from concurrent.futures import ThreadPoolExecutor

def extract_leaf(image):
    # Define the extended range for the entire leaf in the HSV color space
    lower_green = np.array([30, 50, 50])
    upper_green = np.array([90, 255, 255])
    lower_brown = np.array([0, 20, 7])
    upper_brown = np.array([40, 94, 30])

    lower_leaf = np.array([min(lower_green[0], lower_brown[0]),
                           min(lower_green[1], lower_brown[1]),
                           min(lower_green[2], lower_brown[2])])
```

```
    upper_leaf = np.array([max(upper_green[0], upper_brown[0]),
                        max(upper_green[1], upper_brown[1]),
                        max(upper_green[2], upper_brown[2])])

    hsv = cv2.cvtColor(image, cv2.COLOR_BGR2HSV)
    mask = cv2.inRange(hsv, lower_leaf, upper_leaf)


# Extracts the largest object/leaf by selecting the largest contour based on area.

    contours, _ = cv2.findContours(mask, cv2.RETR_EXTERNAL, cv2.CHAIN_APPROX_SIMPLE)

    if contours:
        max_contour = max(contours, key=cv2.contourArea)
        mask = np.zeros(image.shape[:2], dtype=np.uint8)
        cv2.drawContours(mask, [max_contour], -1, (255), thickness=cv2.FILLED)
        leaf_with_transparent_bg = cv2.merge((image[:,:,0], image[:,:,1], image[:,:,2], mask))
        leaf_with_transparent_bg[mask == 0] = [0, 0, 0, 0]
        return leaf_with_transparent_bg
    else:
        return None

# Directories
input_folder = 'input_images/'
output_folder = 'output_images_resized/'

os.makedirs(output_folder, exist_ok=True)

# Creation Excel file
wb = openpyxl.Workbook()
ws = wb.active
ws.append(['Original_image', 'Extraction_successful'])

def process_image(image_path):
    image = cv2.imread(image_path)
    leaf = extract_leaf(image)

    filename = os.path.basename(image_path)
    output_filename = os.path.splitext(filename)[0] + '_leaf.jpg'

    if leaf is not None:
        # Resize to 1856 x 2784
        resized_leaf = cv2.resize(leaf, (1856, 2784))
        output_path = os.path.join(output_folder, output_filename)
        cv2.imwrite(output_path, resized_leaf, [int(cv2.IMWRITE_JPEG_QUALITY), 40])
        ws.append([filename, 'Yes'])
        print(output_filename + " saved")
    else:
        ws.append([filename, 'No'])
        print(output_filename + " not saved")

if __name__ == '__main__':
    with ThreadPoolExecutor() as executor:
        executor.map(process_image, glob.glob(os.path.join(input_folder, '*.JPG')))
```

```
    excel_file = 'Images_info.xlsx'
    wb.save(excel_file)
    print("Excel file created", excel_file)

    print("Process completed.")
```

**Supplementary script 3. Leaf_color_detector.py**

```
""" @author: Matthias Pfeifer """


import os
import cv2
import numpy as np
import pandas as pd

def remove_black_background(image):
    hsv = cv2.cvtColor(image, cv2.COLOR_BGR2HSV)
    lower_black = np.array([0, 0, 0])
    upper_black = np.array([180, 255, 30])
    mask_black = cv2.inRange(hsv, lower_black, upper_black)
    mask_black = cv2.bitwise_not(mask_black)
    result = cv2.bitwise_and(image, image, mask=mask_black)
    return result, mask_black

def calculate_percentage(mask, removed_bg_mask):
    total_pixels = mask.size
    bg_pixels = np.sum(removed_bg_mask == 0)
    valid_pixels = total_pixels - bg_pixels
    mask_pixels = np.sum(mask == 255)
    percentage = (mask_pixels / valid_pixels) * 100 if valid_pixels > 0 else 0
    return percentage, valid_pixels

def apply_morphological_operations(mask):
    kernel = np.ones((5, 5), np.uint8)
    mask_closed = cv2.morphologyEx(mask, cv2.MORPH_CLOSE, kernel)
    return mask_closed

def process_image(image_path):
    image = cv2.imread(image_path)
    image_no_bg, removed_bg_mask = remove_black_background(image)
    hsv = cv2.cvtColor(image_no_bg, cv2.COLOR_BGR2HSV)

    # Adjust color range if needed
    lower_green = np.array([35, 40, 40])
    upper_green = np.array([85, 255, 255])
    lower_yellow = np.array([20, 100, 85])
    upper_yellow = np.array([30, 255, 255])
    lower_brown = np.array([0, 20, 10])
    upper_brown = np.array([30, 255, 170])

    # Masks for green, brown, yellow
```

```
    mask_green = cv2.inRange(hsv, lower_green, upper_green)
    mask_brown = cv2.inRange(hsv, lower_brown, upper_brown)
    mask_yellow = cv2.inRange(hsv, lower_yellow, upper_yellow)

    # Adjustment for grown mask
    mask_brown = apply_morphological_operations(mask_brown)

    # Percentage Calculation
    percentage_green, valid_pixels = calculate_percentage(mask_green, removed_bg_mask)
    percentage_brown, _ = calculate_percentage(mask_brown, removed_bg_mask)
    percentage_yellow, _ = calculate_percentage(mask_yellow, removed_bg_mask)

    # Calculation of unassigned pixels
    total_percentage = percentage_green + percentage_brown + percentage_yellow
    unassigned_percentage = 100 - total_percentage

    # Adjusting the values so that the sum equals 100%
    adjustment_factor = (100 / total_percentage) if total_percentage > 0 else 1
    adjusted_green = percentage_green * adjustment_factor
    adjusted_brown = percentage_brown * adjustment_factor
    adjusted_yellow = percentage_yellow * adjustment_factor

    return percentage_green, percentage_yellow, percentage_brown, unassigned_percentage, adjusted_green,
adjusted_yellow, adjusted_brown

# Input and output settings
input_folder = 'output_images_resized'
output_excel_path = 'output_images_color_detection.xlsx'

# Initialize results
results_data = {'Filename': [], 'Percentage Green': [], 'Percentage Yellow': [], 'Percentage Brown': [], 'Unassigned Pixels
(%)': [],
                'Adjusted Green (%)': [], 'Adjusted Yellow (%)': [], 'Adjusted Brown (%)': []}

for image_file in os.listdir(input_folder):
    if image_file.lower().endswith('.jpg'):
        image_path = os.path.join(input_folder, image_file)

        # Analyze image
        (percentage_green, percentage_yellow, percentage_brown,
         unassigned_percentage, adjusted_green, adjusted_yellow, adjusted_brown) = process_image(image_path)
        results_data['Filename'].append(image_file)
        results_data['Percentage Green'].append(percentage_green)
        results_data['Percentage Yellow'].append(percentage_yellow)
        results_data['Percentage Brown'].append(percentage_brown)
        results_data['Unassigned Pixels (%)'].append(unassigned_percentage)
        results_data['Adjusted Green (%)'].append(adjusted_green)
        results_data['Adjusted Yellow (%)'].append(adjusted_yellow)
        results_data['Adjusted Brown (%)'].append(adjusted_brown)

# Save results to an Excel file
df_results = pd.DataFrame(results_data)
df_results.to_excel(output_excel_path, index=False)
print("Analysis results with adjustments have been saved in", output_excel_path)
```
